# PARP inhibitor synthetic lethality reveals homologous recombination sub-pathway architecture

**DOI:** 10.64898/2026.03.06.709881

**Authors:** Ki Choi Chan, Anastasiia Kovina, Johanna Ertl da Costa, Anke Busch, Ratna N. Cordoni, Björn Stratenwerth, Markus Löbrich

## Abstract

The DNA damage response (DDR) is a complex network of interconnected pathways and sub-pathways that safeguards genome integrity. Deciphering the coordinated and complementary interactions among these pathways remains a major challenge. In this study, we employed CRISPR screening to systematically map the genetic interactions required for different sub-pathways of homologous recombination in human cells following PARP inhibitor treatment. Our approach recapitulated known interactions and uncovered several previously unrecognized connections. We identified RAD54L, in addition to ATRX, as a factor promoting the double Holliday junction (dHJ) pathway and demonstrated that RAD51AP1 and RAD54B function in synthesis-dependent strand annealing (SDSA). We provide evidence that loss of TOP3A induces a switch in HR sub-pathway usage from SDSA to the dHJ pathway. Furthermore, TOP3A deficiency abolishes the requirement for ATRX and the histone variant H3.3 in the dHJ pathway, while maintaining strict dependence on RAD54L. We further observed that H3.3 is involved in both HR sub-pathways, whereas its depositing chromatin remodelers HIRA and ATRX play pathway-specific roles in SDSA and dHJ, respectively. Together, our findings define the architecture underlying HR sub-pathway choice and reveal a key role for TOP3A in regulating pathway balance.

## Introduction

The DNA damage response (DDR) is a highly coordinated signalling and repair network that preserves genome stability in response to endogenous and exogenous genotoxic stress. DNA double-strand breaks (DSBs) represent highly cytotoxic lesions that jeopardize genomic stability and cell survival. To counteract them, cells employ multiple repair pathways, whose engagement is dictated by cell type, cell-cycle stage, and local chromatin or sequence context^1^. Canonical non-homologous end joining (c-NHEJ), in particular alternative end joining (alt-EJ) and single-strand annealing (SSA) operate in an error-prone manner and frequently generate genomic rearrangements. In contrast, homologous recombination (HR) provides a high-fidelity repair mechanism that preserves genome stability during DSB repair.

HR is an evolutionarily conserved pathway that uses an intact homologous DNA sequence, most commonly the sister chromatid, as a template to accurately restore genetic information at the site of damage^2–3^. Consequently, HR is largely restricted to the S and G2 phases of the cell cycle, when a sister chromatid is available. HR proceeds through a series of coordinated steps. It is initiated by nucleolytic processing of DNA ends (resection), which generates extended 3′ single-stranded DNA (ssDNA) overhangs that are subsequently coated by the ssDNA-binding protein RPA. BRCA2 facilitates the replacement of RPA with RAD51, promoting assembly of the presynaptic filament that mediates homology search and pairs with a complementary DNA template. Successful homology recognition triggers strand invasion and the formation of a displacement loop (D-loop)^4^. This initial intermediate, termed the nascent D-loop, transitions to an extending D-loop upon DNA polymerase-mediated DNA synthesis. The structure of D-loops and the way extending D-loops are processed is a key determinant of HR sub-pathway usage and repair outcome. Repair of two-ended DSBs by HR can occur through two distinct sub-pathways. One pathway involves the formation of a double Holliday junction (dHJ) structure following D-loop formation and extended DNA repair synthesis, with the potential to generate crossover events between sister chromatids. In contrast, synthesis-dependent strand annealing (SDSA) is associated with short DNA repair synthesis patches and involves displacement of the invading strand after repair synthesis and annealing to the second break end, thereby avoiding dHJ formation and crossover outcomes. These two sub-pathways can compensate for one another to ensure efficient repair.

Early efforts to elucidate the molecular mechanisms of the DDR focused largely on individual repair proteins and pathways. However, nowadays it is clear that DDR pathways are highly interconnected, with extensive functional overlap that allows them to compensate for each other. This complexity has created a need for experimental approaches capable of systematically mapping combinatorial genetic interactions within the DDR. Genome-wide CRISPR knockout screens have emerged as powerful tools to address this challenge, enabling investigation of interactions between pathways required for successful DSB repair and cellular survival^5–8^. These approaches exploit synthetic lethality, which means that cells tolerate the loss of single genes but not their combined inactivation, thereby revealing functional redundancy and alternative compensatory mechanisms within the DDR.

The earliest CRISPR–Cas9 knockout screens established the synthetic lethal relationship between BRCA1/2 and PARP^9^. Subsequent application of this approach uncovered key modulators of PARP inhibitor (PARPi) response^9–11^, as well as additional synthetically lethal partners of BRCA1/2. For example, genome-wide CRISPR knockout (KO) screens performed in BRCA1-deficient cells treated with PARP inhibitors or other genotoxic agents identified 53BP1 and its downstream effectors (RIF1, Shieldin) as top resistance hits, as their loss restores DNA end resection and confers resistance to PARP inhibition^12–14^. Conversely, negative selection hits, whose loss further sensitizes HR-deficient cells, revealed numerous additional factors, including the DNA endonuclease APE2, flap endonuclease 1 (FEN1), polymerase θ (POLQ), and the protein phosphatase 2A inhibitor CIP2A^15–19^. These discoveries have provided a rich resource for understanding compensatory DNA repair pathways in cells lacking HR and have opened new opportunities for targeting cancer cells by exploiting their reliance on backup DNA damage response mechanisms.

Recent advances have improved CRISPR-based screens using DDR-focused libraries that enhance sensitivity and depth of analysis^20–22^. Additionally, over the past years, combinatorial CRISPR knockout screens, using dual-gRNA libraries focused on DNA damage response genes, have been developed to systematically enable simultaneous perturbation of gene pairs, allowing high-resolution mapping of genetic interactions and pathway overlap within the DDR^23–25^. Early dual-guide DDR screens revealed that a p53-mediated response reduces the sensitivity of CRISPR–Cas9 screens^20^. Moreover, such screens have uncovered several previously unrecognized DDR functional interactions. Notable examples include the synthetic lethality between the WDR48–USP1 deubiquitinating complex and single-strand break repair enzymes FEN1 or LIG1, where loss of both activities leads to DNA gap accumulation, aberrant PCNA ubiquitylation, replication failure, and genome instability^26^. This highlights USP1 as a potential therapeutic target in FEN1-mutant cancers. Another example is the synthetic lethality between the translocases FANCM and SMARCAL1, whose combined loss results in unresolved replication-associated DNA structures and accumulation of DSBs that persist into mitosis, suggesting therapeutic vulnerabilities in tumors lacking either factor. In other studies, combinatorial DDR CRISPR KO screens identified synthetically lethal interactions under both basal conditions and following ionizing irradiation, which predominantly induces DSBs, yielding hits associated with either sensitivity or resistance^27^. However, this study also highlights several limitations of dual-gRNA approaches, including high recombination rates that lead to the loss of a substantial fraction of sequencing reads, as well as the inability to model gradual mutational adaptation in cancer cells, as dual guide RNA combinatorial screens induce simultaneous knockout of two genes, potentially explaining the inability to find some known genetic interactions.

Because of the functional redundancy of HR sub-pathways, we conducted a systematic analysis of the genetic interactions that regulate homologous recombination. We applied a three-dimensional CRISPR screening approach across three cell lines, enabling parallel comparison of their genetic dependencies. This strategy allowed detailed mapping of HR sub-pathways and their associated factors, including functional annotation of sub-pathway-specific genes. We confirmed RAD54L as the dHJ pathway-promoting factor and showed RAD51AP1 and RAD54B as mediators of SDSA. Moreover, we demonstrated that TOP3A loss shifts repair from SDSA to dHJ while bypassing ATRX but retaining RAD54L dependence. Notably, the histone variant H3.3 contributes to both HR sub-pathways, whereas its chromatin remodelers HIRA and ATRX act in a pathway-specific manner.

## Results

### A 3-D CRISPR Cas9 dropout screen identifies distinct networks of HR factors

To explore the networks of HR factors underlying PARPi sensitivities, we made use of the described synthetic lethality between RAD54L and RAD51AP1 (henceforth abbreviated as 54L and AP1, respectively)^28^. We wished to identify the network of HR factors associated with the functions of 54L and AP1 and performed a 3-D CRISPR Cas9 KO dropout screen. We generated HeLa 54L and AP1 KO cells (Fig. S1A) and first investigated cell viability after treatment with the PARPi olaparib using siRNA-mediated gene depletion. The combined depletion of 54L and AP1 sensitized HeLa empty vector (EV) cells to PARPi, similar to BRCA2 depletion, whereas the single depletion of 54L or AP1 had no effect (Fig. S1A). In contrast, the single depletion of AP1 in 54L KO cells and the single depletion of 54L in AP1 KO cells phenocopied the PARPi sensitivity of siBRCA2 (Fig. S1A). EV cells are HeLa wild-type (WT) cells transfected with a Cas9-expressing plasmid without sgRNA.

We then performed a genome-wide CRISPR-Cas9 screen to identify genes whose loss leads to PARPi sensitivity in the three cell lines. The Brunello library was used for this approach, which harbors 76,440 short guide RNAs (sgRNAs) targeting 19,000 genes^47^. We transduced the cell lines with the library at a low multiplicity of infection (MOI) which allowed co-expression of Cas9 along with the sgRNAs and results in only one integrant per cell. After antibiotic selection, we grew the cells with or without PARPi treatment. Genomic DNA was harvested for library preparation, followed by NGS and bioinformatic analysis to quantify the sgRNAs present in the remaining populations (Fig. S1B). We created a bioinformatics pipeline based on the established MaGECK algorithm to identify both positively and negatively selected genes in the three cell lines (Fig. S1C).

We plotted the β-scores in three panels corresponding to the three combinations of a pairwise comparison between the cell lines (Fig. 1A). The plots identify PARP1 and PARG as positive hits in all three lines, confirming that loss of these factors mediate PARPi resistance. As expected, we found 54L as a strong negative hit in the AP1 KO line and AP1 as a strong negative hit in the 54L KO line, while loss of 54L and AP1 had no effect in their corresponding KO lines (see Fig. 1Ai for the comparison between the AP1 and 54L KO lines). Interestingly, 54L and AP1 were also identified as hits in HeLa EV cells, suggesting that the single loss of these factors causes PARPi sensitivity in this experiment (Fig. 1Aii and 1Aiii). Moreover, factors known to be essential for HR, such as BRCA1 and BARD1, were identified as hits in all three lines, further confirming the validity of the screen (Fig. 1A).

**Figure 1.**
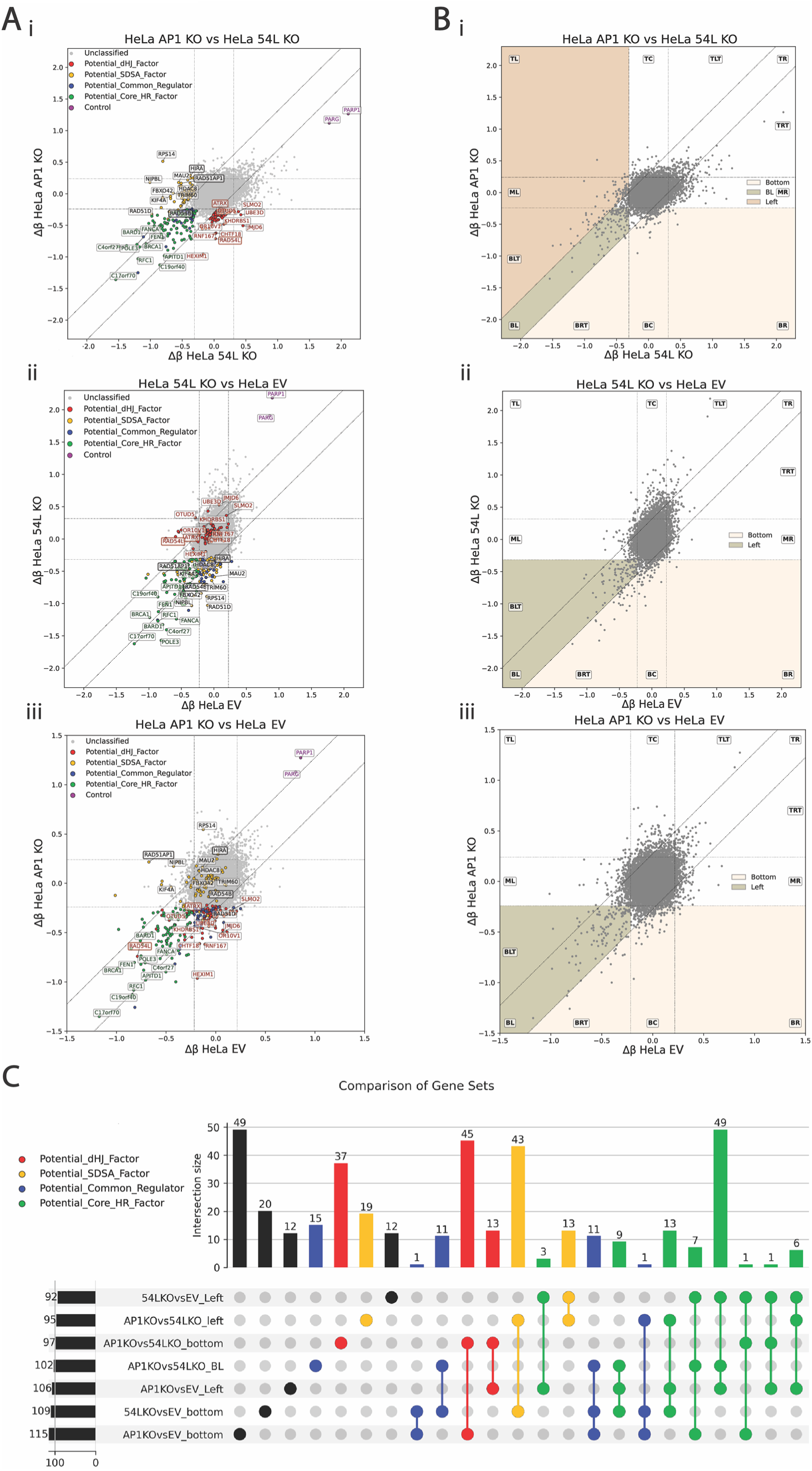
CRISPR screens identify synthetic lethal targets in HeLa EV, RAD54L KO and RAD51AP1 KO cells. **(A)** Scatter plots of gene-level Δβ-scores comparing (i) HeLa AP1 KO vs HeLa 54L KO, (ii) HeLa 54L KO vs HeLa EV and (iii) HeLa AP1 KO vs HeLa EV. For each gene, the Δβ-score was calculated per cell line as the difference between PARPi-treated and untreated (NT) β-scores (Δβ = β_PARPi_ - β_NT_). Each point represents one gene plotted by the Δβ-scores of the two compared cell lines (x-axis and y-axis), where more negative Δβ-scores indicate stronger sensitization. Cutoff lines were defined using the standard deviation (σ) of the respective Δβ distributions: horizontal cutoffs correspond to ±2σ of the y-values, vertical cutoffs correspond to ±2σ of the x-values, and diagonal cutoffs correspond to ±2σ of the difference between the two cell lines’ Δβ-scores (Δβ_y_ - Δβ_x_). Genes assigned to the UpSet-derived hit groups (Figure 1C) are highlighted in the corresponding colour code, while non-highlighted genes are shown in grey. The top candidate genes are annotated (top 10 genes for potential dHJ-factors, potential SDSA-factors and potential Core-HR-factors). Label colours match the assigned hit group; for readability, labels for potential SDSA-factors are shown in black. Selected genes of interest are additionally emphasized by a thicker label border. Controls (PARP1 and PARG) are indicated in purple. **(B)** Scatter plots comparing Δβ-scores between genetic backgrounds (i) HeLa AP1 KO vs HeLa 54L KO, (ii) HeLa 54L KO vs HeLa EV and (iii) HeLa AP1 KO vs HeLa EV. Cutoff lines and region definitions are identical to Figure 1A. Shaded regions indicate gene sets extracted for downstream overlap analysis; sgRNAs that preferentially sensitize one knockout background to olaparib were selected. Abbreviations: TL, top left; TC, top center; TLT, top left triangle; TR, top right; TRT, top right triangle; MR, middle right; BR, bottom right; BC, bottom center; BRT, bottom right triangle; BL, bottom left; BLT, bottom left triangle; ML, middle left. **(C)** UpSet plot summarizing overlap between gene sets derived from Figure 1B. Horizontal bars represent the number of genes in each defined region, vertical bars indicate intersection size, and the dot matrix indicates which gene sets contribute to each intersection. Intersections are color-coded as follows: red, potential dHJ-factors; yellow, potential SDSA-factors; blue, potential Common Regulators; green, potential Core-HR-factors.

To systematically explore the network of factors which are either involved specifically with the functions of 54L or AP1, or more generally promote HR, we divided the 2-D β-score plots of the three combinations of cell lines in figure 1A into different areas (Fig. 1B). For the comparison between the AP1 KO with the 54L KO cell line, we grouped factors that only sensitize the AP1 KO line (area “left”), only the 54L KO line (area “bottom”) or both cell lines similarly (area “bottom left”). For the comparison between the AP1 KO or the 54L KO cell line with the EV line, we grouped factors that only sensitize the AP1 KO or the 54L KO line but not the EV line (area “bottom”) or both AP1 KO or 54L KO and the EV line (area “left”). To simplify the different combinations of areas for the three different plots, we generated an UpSet diagram (advanced Venn diagram) showing all combinations of categories together with the number of hits found and our assignment as specific AP1 or 54L factors or as general HR factors (Fig. 1C).

We reasoned that factors specifically associated with the function of 54L should fall into the “bottom” category of the AP1 KO vs. 54L KO plot and either into the “left” or “bottom” category of the AP1 KO vs. EV plot, depending on whether or not their loss also sensitizes EV cells. However, if they additionally fall into the “left” or “bottom” category of the 54L KO vs. EV plot, they should be considered general or core HR factors since their loss also sensitizes 54L KO cells. Accordingly, we expected factors specifically associated with the function of AP1 to fall into the “left” category of the AP1 KO vs. 54L KO plot and either into the “left” or the “bottom” category of the AP1 KO vs. EV plot, depending on whether or not their loss also sensitizes EV cells, but not into the ‘left’ or “bottom” category of the AP1 KO vs. EV plot. Moreover, additional factors which promote HR in general should fall into the “bottom left” category of the AP1 KO vs. 54L KO plot and should be in the “left” category in either the AP1 KO vs. EV or the 54L KO vs. EV plot, as we reasoned that loss of core HR factors should sensitize EV cells more than at least one of the two KO cell lines. A more detailed discussion of our factor assignment is provided in the supplement.

Most 54L-associated factors fall only into the category “AP1 KO vs. 54L KO bottom” or simultaneously into this category and into the category “AP1 KO vs. EV bottom” (and less frequently into the “AP1 KO vs. 54L KO bottom” and “AP1 KO vs. EV left” categories). Similarly, most AP1-associated factors fall only into the category “AP1 KO vs. 54L KO left” or into this category and into the category “54L KO vs. EV bottom” (and less frequently into the “AP1 KO vs. 54L KO left” and “54L KO vs. EV left” categories). This analysis shows that most, but not all, factors which are specifically associated with the functions of either AP1 or 54L are dispensable for PARPi resistance of EV cells (Fig. 1C). Most general HR factors fall into the “bottom left” category of the AP1 KO vs. 54L KO plot and into the “left” category of both the AP1 KO vs. EV and the 54L KO vs. EV plots, indicating that most core HR factors sensitize EV cells more than both KO cell lines. In all, our 3-D CRISPR Cas9 screen identified 95 factors that are specifically associated with the function of 54L, 75 factors specifically associated with the function of AP1, and 89 factors that are important for PARPi resistance in all three cell lines. A list of these factors is provided in Table S1.

### RAD54L, RAD51AP1 and RAD54B promote distinct HR sub-pathways

We noted from our screening results that ATRX, a factor which we previously identified to promote DSB repair by the dHJ HR sub-pathway^29–30^, is specifically associated with the function of 54L, while RAD54B (54B), a paralog of 54L, is specifically associated with the function of AP1. This finding prompted us to investigate if the two HR networks associated by the functions of 54L and AP1 might represent the dHJ and the SDSA HR sub-pathway, respectively.

First, we confirmed that the loss of 54L sensitizes 54B KO but not EV cells to PARPi treatment (Fig. S2A). We then studied the repair of DSBs arising from PARPi treatment. For this, we treated HeLa cells with PARPi for 24 h and scored γH2AX foci as a proxi for unrepaired DSBs in G2-phase cells (Fig. 2A). We previously showed that DSBs after PARPi treatment arise during S-phase and are repaired in G2 in a manner dependent on BRCA2^31^. Here, we observed that the single siRNA-mediated depletions of AP1, 54B or 54L did not result in significantly elevated γH2AX foci levels. However, the combined depletions of 54L and AP1, of 54L and 54B, but not of AP1 and 54B, caused γH2AX foci levels that were elevated similar to the repair defect observed after depleting BRCA2 (Fig. 2A-B). We also analyzed AP1, 54L and 54B KO cells treated with siRNA against AP1, 54L or 54B and confirmed that the dual loss of 54L and AP1 and of 54L and 54B but not of AP1 and 54B caused elevated γH2AX foci levels (Fig. S2B). These repair studies confirm that AP1 and 54B operate in the same HR sub-pathway for DSB repair, which is distinct and largely redundant to the sub-pathway promoted by 54L. Moreover, the data also suggest that a failure to repair DSBs underlies the PARPi sensitivities assessed in our CRISPR-Cas9 screen. Lastly, we investigated the effects of combined loss of RAD54L, RAD51AP1, or RAD54B in a HeLa pGC HR reporter cell line^32^. DSB induction by the expression of the I-SceI endonuclease followed by HR repair of the DSB yields GFP-positive cells. We observed that the single loss of 54L or AP1 only partially reduced the level of GFP-positive cells while the combined loss nearly completely abolishes HR (Fig. S2C). The depletion of 54B also further reduced the GFP level of 54L-depleted cells but not of AP1-depleted cells, confirming that 54B operates in the same HR sub-pathway as AP1 (Fig. S2C).

**Figure 2.**
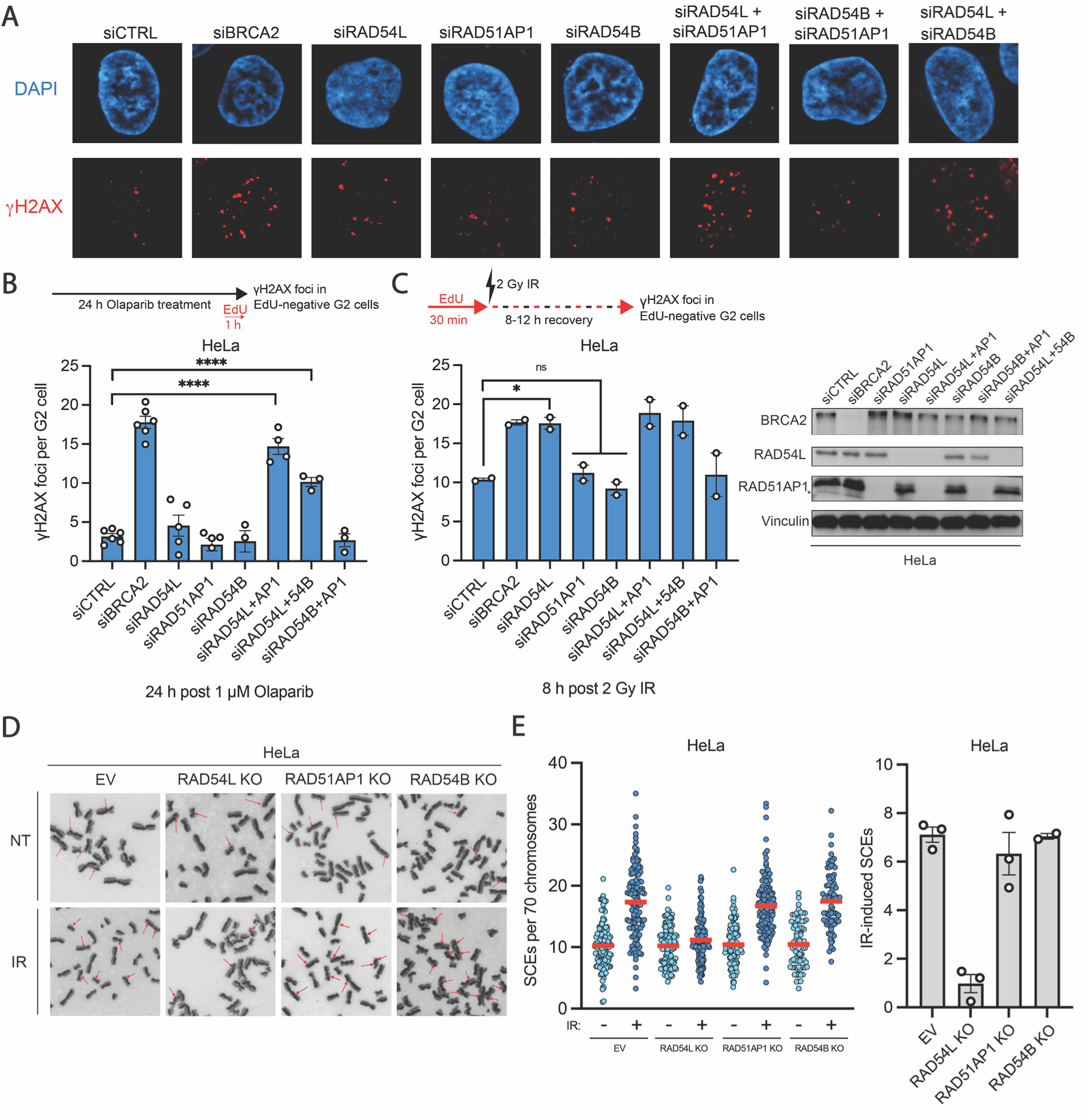
RAD51AP1/RAD54B and RAD54L define distinct sub-pathways of HR. **(A)** Representative images of γH2AX foci in G2 HeLa cells following transfection with the indicated siRNAs and subsequent PARPi treatment are shown. **(B)** (Top) Schematic illustration depicting the treatment strategy for PARPi treatment: cells were treated with 1 µM olaparib for 24 h. 1 h before fixation, 10 µM EdU were added. (Bottom) HeLa cells were transfected with the indicated siRNAs, and γH2AX foci were enumerated after PARPi treatment for 24 h in EdU-negative G2 cells. Spontaneous foci were subtracted. All data show mean ± SEM (n = 3-6). Results from individual experiments, each derived from 40 cells, are indicated. **(C)** (Top) Schematic illustration depicting the irradiation strategy: 10 µM EdU were added 30 min before irradiation with 2 Gy and left on during the recovery period (8 to 12 h). (Bottom) HeLa cells were transfected with the indicated siRNAs, and γH2AX foci were enumerated 8 h following 2 Gy IR in EdU-negative G2 cells. Spontaneous foci were subtracted. All data show mean ± SEM (n = 3-5). Results from individual experiments, each derived from 40 cells, are indicated. Knockdown of BRCA2, RAD54L and RAD51AP1 was confirmed by western blotting (right). Knockdown of RAD54B was confirmed by qPCR (Fig. S2E). **(D)** HeLa WT, HeLa RAD54L KO, HeLa RAD51AP1 KO, and HeLa RAD54B KO cells were irradiated with 2 Gy, collected after 8 h, and processed to obtain mitotic spreads. Representative images of chromosome spreads of nonirradiated and irradiated cells are shown. Red arrows show individual SCE events. **(E)** SCEs per spread were quantified and normalized to 70 chromosomes. Results from individual experiments, each derived from 40 cells, are indicated, and red horizontal lines indicate the mean (left). The bar graph shows the IR-induced SCE numbers (total SCEs after background subtraction) (right). IR-induced SCE data show mean ± SEM, and results from individual experiments are indicated. 120 spreads and >7,000 chromosomes per condition were analysed from three independent experiments. *P < 0.05; **P < 0.01; ***P < 0.001; ****P < 0.0001; ns: not significant (two-tailed t test).

We next investigated DSB repair by HR in U2OS cells devoid of ATRX. We treated U2OS cells with PARPi for 24 h and scored γH2AX foci in G2-phase cells. In contrast to HeLa cells, the single depletion of AP1 or 54B caused elevated foci numbers, while depletion of 54L, either alone or in combination with AP1 or 54B, did not cause a major defect (Fig. S2D). We also investigated U2OS DR-GFP cells carrying a GFP-based HR reporter system^33^ and found that depletion of RAD54L alone did not affect the proportion of GFP-positive cells (Fig. S2F). In contrast, single depletion of RAD51AP1 or RAD54B partially reduced GFP levels, and co-depletion of RAD54L with either factor led to a further decrease in GFP signal. In conclusion, this analysis suggests that 54L is largely dispensable in cells devoid of ATRX and reinforces the notion that 54L and ATRX operate in the same HR sub-pathway. In contrast to U2OS cells, ATRX-positive HeLa cells employ both HR sub-pathways for repairing PARPi-induced DSBs.

To more specifically define the HR sub-pathways promoted by AP1 and 54B on one hand and by 54L and ATRX on the other hand, we investigated DSB repair by HR in G2-irradiated cells. We have previously used this approach to show that ATRX promotes the formation of sister chromatid exchanges (SCEs) arising from DSB repair by the dHJ sub-pathway^29–30^. Since SCEs can also arise at stalled replication forks or during the repair of post-replicative gaps, SCE formation from S-phase-damaging agents such as PARPi cannot be used to evaluate HR sub-pathway usage^34^. However, after irradiation of G2-phase cells, SCEs exclusively arise from DSB repair by the dHJ pathway^35^. We first assessed the level of unrepaired DSBs by scoring γH2AX foci in irradiated G2-phase cells as previously shown^29–30^. Consistent with previous observations, depletion of 54L in HeLa cells caused elevated foci numbers similar to the depletion of BRCA2^36^ (Fig. 2C). In contrast, depletion of AP1 or 54B had only a minor impact (Fig. 2C). We then measured the formation of SCEs in G2-irradiated HeLa cells and observed strongly reduced SCE formation in cells deficient for 54L but not for AP1 or 54B (Fig. 2D-E). This finding is consistent with our previous notion that ATRX-proficient cells predominantly use the dHJ sub-pathway for repairing DSBs in G2^30^. In contrast to HeLa cells, depletion of 54L did not lead to elevated foci numbers in G2-irradiated U2OS cells, while the depletion of AP1 or 54B caused a repair defect similar to the depletion of BRCA2 (Fig. S3A). Since we previously established that HR in G2-irradiated U2OS cells proceeds by SDSA without SCE formation^30^, this result supports the notion that while 54L and ATRX promote the dHJ sub-pathway, AP1 and 54B promote SDSA.

To confirm and extend these findings, we employed U2OS^ATRX^ cells which express ATRX upon doxycycline (Dox) treatment^37^, allowing the investigation of HR sub-pathway usage in dependency of ATRX in the same cell system. Without Dox treatment, DSB repair in G2-irradiated U2OS^ATRX^ cells depended on AP1 and 54B, whereas 54L was dispensable (Fig. 3A-B). However, after the expression of ATRX by Dox treatment, 54L became necessary, while AP1 and 54B were dispensable (Fig. 3A-B), recapitulating the results with ATRX-proficient HeLa cells. We then assessed SCE formation in G2-irradiated U2OS^ATRX^ cells, which was induced by the expression of ATRX (Fig. 3C). Depletion of 54L but not AP1 or 54B completely abolished SCE formation in U2OS^ATRX^ cells expressing ATRX (Fig. 3C). This result confirms our analysis with HeLa cells and establishes 54L in addition to ATRX as a dHJ factor and AP1 as well as 54B as SDSA factors (Fig. 3D).

**Figure 3.**
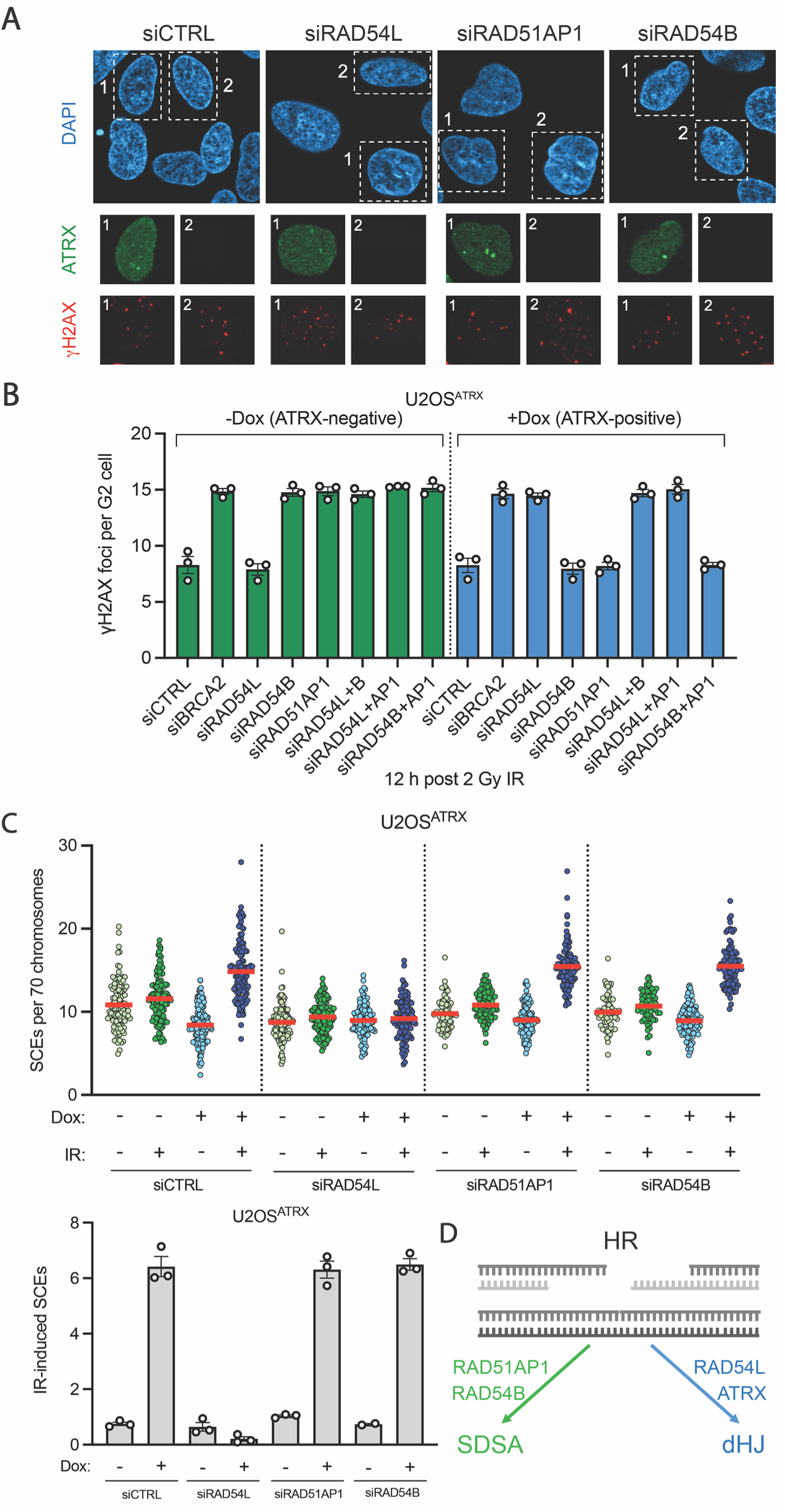
ATRX induction triggers a switch from RAD51AP1/RAD54B-dependent SDSA to a RAD54L-dependent dHJ sub-pathway. **(A)** Representative images of γH2AX foci and ATRX expression in U2OS^ATRX^ G2 cells following transfection with the indicated siRNAs and subsequent irradiation are shown. **(B)** U2OS^ATRX^ cells, with (blue) and without (green) doxycycline-induced ATRX expression, were transfected with the indicated siRNAs, and γH2AX foci were enumerated at 12 h following 2 Gy IR in EdU-negative G2 cells. Spontaneous foci were subtracted. All data show mean ± SEM (n = 3). Results from individual experiments, each derived from 40 cells, are indicated. Knockdown was confirmed by western blotting. **(C)** U2OS^ATRX^ cells, with (blue) and without (green) doxycycline-induced ATRX expression, were transfected with the indicated siRNAs. They were irradiated with 2 Gy, collected after 16h, and processed to obtain chromosome spreads. SCEs per spread were quantified and normalized to 70 chromosomes. Results from individual experiments, each derived from 40 cells, are indicated (top), and red horizontal lines indicate the mean. The bar graph shows the IR-induced SCE numbers (total SCEs after background subtraction) (bottom). IR-induced SCE data show mean ± SEM, and results from individual experiments are indicated. 80-120 spreads and >7,000 chromosomes per condition were analysed from 2-3 independent experiments. **(D)** Model illustrating the interplay between RAD51AP1 and RAD54B, which promote SDSA, and RAD54L and ATRX, which facilitate dHJ formation.

### TOP3A regulates HR sub-pathway choice

Since 54L, AP1 and 54B have been shown to operate at the level of D-loop formation, we next studied if and how D-loop formation can regulate HR sub-pathway choice. Studies in yeast have suggested that Rad54 and Rdh54 (likely homologues of human 54L and 54B, respectively) promote the formation of two distinct types of D-loops, which are differentially regulated by Top3^38–39^. Therefore, we investigated the effect of human TOP3A on HR sub-pathway regulation in U2OS and HeLa cells. First, we depleted TOP3A by siRNA and assessed unrepaired γH2AX foci levels in G2-irradiated U2OS cells. The depletion of TOP3A did not affect foci levels in WT cells but reduced the elevated γH2AX foci level of AP1- or 54B-deficient cells to that of WT cells (Fig. S4A). It also rescued the elevated γH2AX foci level of cells deficient for RECQ5, a factor which we previously showed to be involved in SDSA^30^, suggesting that TOP3A-depleted U2OS cells utilize a repair pathway different to SDSA (Fig. S4A). Similar results were obtained after the depletion of BLM, which has been shown to function in a complex with TOP3A during D-loop regulation^40^ (Fig. S4E). To explore if repair in the absence of TOP3A/BLM still represents HR, we assessed RAD51 foci levels at 2 h post IR, a time at which RAD51 foci peak in irradiated G2 cells^30^. Depletion of TOP3A did not affect RAD51 foci levels (Fig. S4D), suggesting that U2OS cells devoid of TOP3A utilize an HR pathway. Strikingly, we observed that the depletion of 54L, which by itself did not affect γH2AX foci levels in G2-irradiated U2OS cells, caused elevated foci levels if combined with the depletion of TOP3A (Fig. S4A). We also observed that the depletion of TOP3A rescues the elevated γH2AX foci level of AP1- or 54B-deficient U2OS cells after PARPi treatment while causing elevated foci numbers in 54L-deficient cells, showing that the switch in the requirement from AP1/54B to 54L is not a peculiarity of irradiation (Fig. S4B). Finally, we observed that the HR deficiency of U2OS cells deficient for AP1, 54B or RECQ5 in a reporter assay is also rescued by the depletion of TOP3A (Fig. S4C). Together, these findings suggest that U2OS cells devoid of TOP3A utilize a 54L-dependent repair process, which, in ATRX-proficient HeLa cells, represents the dHJ sub-pathway.

To investigate the possibility that ATRX-deficient U2OS cells employ the dHJ pathway, we used the U2OS^ATRX^ cells. We first showed that without Dox addition the depletion of TOP3A reduced the IR-induced elevated γH2AX foci levels of AP1- and 54B-depleted cells to the level of WT cells while causing elevated foci numbers in 54L-deficient cells (Fig. 4A-B), phenocopying the results obtained with normal U2OS cells (Fig. S4A). Interestingly, the expression of ATRX by Dox treatment caused a requirement for 54L, which was not affected by the additional depletion of TOP3A (Fig. 4B). We then assessed SCE formation in G2-irradiated U2OS^ATRX^ cells and observed IR-induced SCE levels after TOP3A-depletion in ATRX-deficient U2OS cells similar to the SCE levels observed in these cells after the expression of ATRX. SCEs induced by either the depletion of TOP3A or by the expression of ATRX were both strictly dependent on 54L, and the depletion of TOP3A combined with the expression of ATRX did not further enhance the IR-induced SCE levels (Fig. 4C). We also assessed the extent of DNA repair synthesis during HR by the analysis of BrdU foci^41^. Our previous studies showed that DNA repair synthesis occurring during the dHJ sub-pathway, but not during SDSA, leads to visible BrdU foci^29–30^. Here, we observed that the depletion of TOP3A in U2OS cells caused IR-induced BrdU foci which were dependent on 54L (Fig. 4D-E). Collectively, these findings strongly suggest that loss of TOP3A function in U2OS cells leads to a switch in HR sub-pathway usage from SDSA to the dHJ pathway (Fig. 4F).

**Figure 4.**
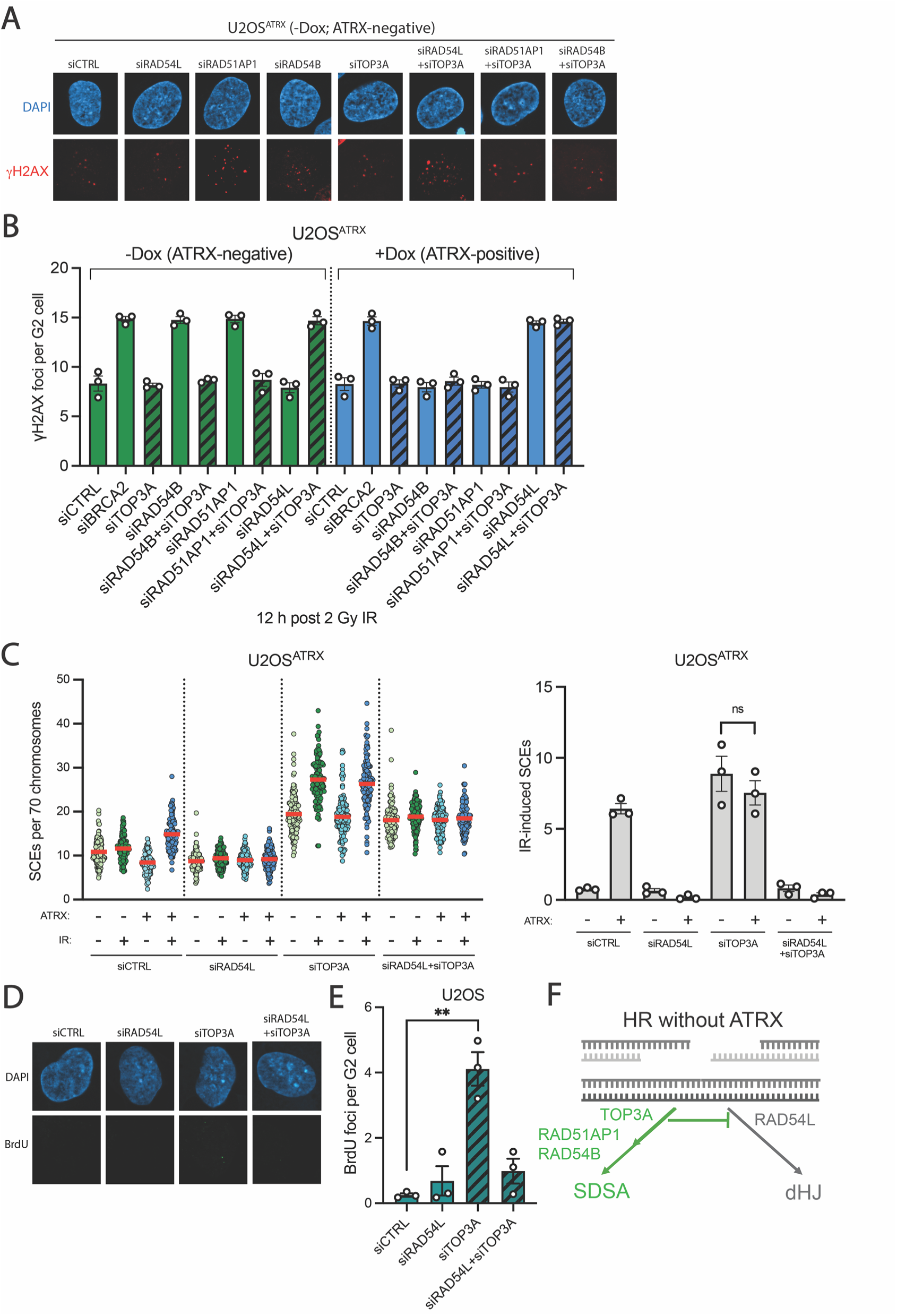
Loss of TOP3A function in ATRX-deficient U2OS cells causes a shift in HR sub-pathway usage from SDSA to the RAD54L-dependent dHJ sub-pathway. **(A)** Representative images of γH2AX foci in U2OS^ATRX^ G2 cells following transfection with the indicated siRNAs and subsequent irradiation are shown. **(B)** U2OS^ATRX^ cells, with (blue) and without (green) doxycycline-induced ATRX expression, were transfected with the indicated siRNAs, and γH2AX foci were enumerated at 12 h following 2 Gy IR in EdU-negative G2 cells. Spontaneous foci were subtracted. Striped bars indicate TOP3A depletion. All data show mean ± SEM (n = 3). Results from individual experiments, each derived from 40 cells, are indicated. Knockdown was confirmed by western blotting. **(C)** U2OS^ATRX^ cells, with (blue) and without (green) doxycycline-induced ATRX expression, were transfected with the indicated siRNAs. They were irradiated with 2 Gy, collected after 16h, and processed to obtain chromosome spreads. SCEs per spread were quantified and normalized to 70 chromosomes. Results from individual experiments, each derived from 40 cells, are indicated (left), and red horizontal lines indicate the mean. The bar graph shows the IR-induced SCE numbers (total SCEs after background subtraction) (right). IR-induced SCE data show mean ± SEM, and results from individual experiments are indicated. 120 spreads and >7,000 chromosomes per condition were analysed from three independent experiments. **(D)** U2OS cells were transfected with siCTRL, siRAD54L and/or siTOP3A, labeled with EdU, irradiated with 4 Gy, and then incubated with BrdU. Representative images of BrdU foci reflecting DNA repair synthesis in G2 cells are shown. **(E)** BrdU foci were enumerated in EdU-negative G2 cells. Spontaneous foci were subtracted. All data show mean ± SEM (n = 3). Results from individual experiments, each derived from 40 cells, are indicated. **P < 0.01 (two-tailed t test). **(F)** Model illustrating the regulation of HR sub-pathway usage by TOP3A in intrinsically ATRX-deficient U2OS cells. TOP3A acts as a positive SDSA-promoting HR factor together with RAD51AP1 and RAD54B to facilitate SDSA usage. Additionally, TOP3A indirectly inhibits the usage of the dHJ sub-pathway in ATRX-deficient cells. *P < 0.05; **P < 0.01; ***P < 0.001; ****P < 0.0001; ns: not significant (two-tailed t test).

### ATRX overcomes the function of TOP3A

Our findings above show that TOP3A functions in ATRX-deficient U2OS cells to overcome a requirement for 54L and causes a dependency on AP1/54B (Fig. 4F). The expression of ATRX in U2OS cells prevents the switch from 54L-dependent to AP1/54B-dependent HR. We therefore reasoned that HeLa cells, which we previously showed to depend on ATRX for HR repair^29–30^, might perform ATRX-independent repair if TOP3A is absent. To test this idea, we depleted ATRX and/or TOP3A in HeLa cells and measured unrepaired γH2AX foci levels in G2-irradiated cells (Fig. 5A). While the single depletion of TOP3A did not cause elevated foci levels, the depletion of ATRX or 54L conferred a repair defect, similar to our previous study^29^ (Fig. 5A). Strikingly, the depletion of TOP3A rescued the elevated foci level of ATRX-depleted but not of 54L-depleted cells (Fig. 5A). ATRX functions during the dHJ pathway by depositing the histone 3.3 (H3.3) to promote DNA repair synthesis and SCE formation^29^. We therefore also depleted H3.3 in combination with TOP3A and observed elevated γH2AX foci levels after the single depletion of H3.3, which were rescued by the additional depletion of TOP3A (Fig. 5A). We also studied ATRX-KO HeLa cells and confirmed that the co-depletion of TOP3A rescues the elevated foci numbers even in the absence of H3.3 (Fig. S5). Furthermore, using the HeLa pGC reporter system, the reduced HR activity after the loss of ATRX or 54L were substantially rescued by co-depletion of TOP3A (Fig. 5B). These results indicate that HR is largely restored in HeLa cells lacking TOP3A and ATRX/H3.3.

**Figure 5.**
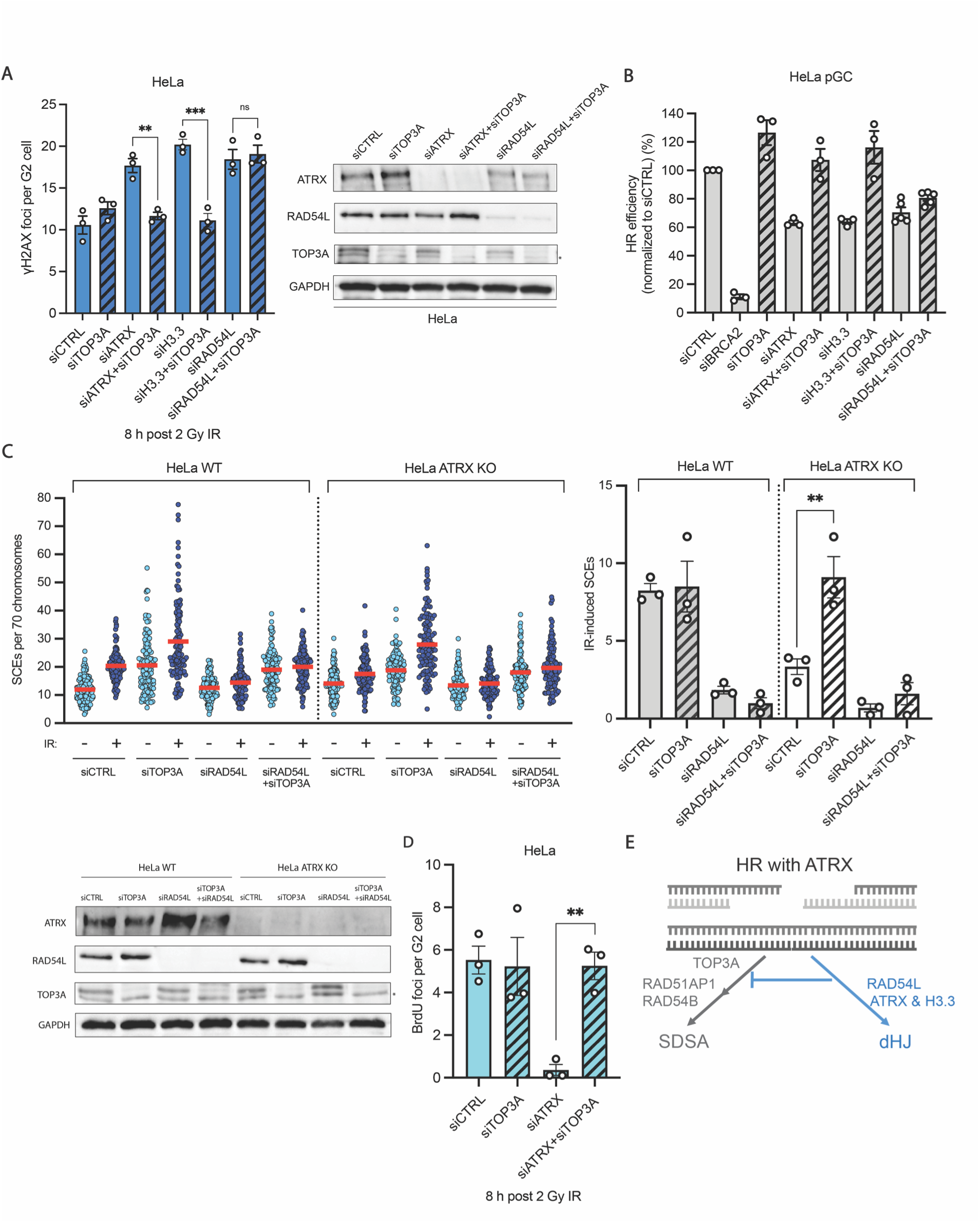
ATRX, together with H3.3, antagonizes TOP3A function and promotes the dHJ pathway in HeLa cells. **(A)** HeLa cells were transfected with the indicated siRNAs, and γH2AX foci were enumerated 8 h following 2 Gy IR in EdU-negative G2 cells. Spontaneous foci were subtracted. All data show mean ± SEM (n = 3-5). Results from individual experiments, each derived from 40 cells, are indicated. Knockdown of TOP3A, ATRX and RAD54L was confirmed by western blotting (right). **(B)** HeLa pGC reporter cells carry a GFP cassette that is disrupted by the I-SceI sequence and a second truncated GFP cassette. I-SceI-induced HR repair events result in the expression of functional GFP. Cells were transfected with the indicated siRNAs followed by the I-SceI plasmid transfection, and GFP-positive cells were measured by flow cytometry. All data show mean ± SEM (n = 3-5). Results from individual experiments are indicated. **(C)** HeLa WT and HeLa ATRX KO cells were irradiated with 2 Gy, collected after 8 h, and processed to obtain mitotic spreads. SCEs per spread were quantified and normalized to 70 chromosomes. Results from individual experiments, each derived from 40 cells, are indicated, and red horizontal lines indicate the mean (left). The bar graph shows the IR-induced SCE numbers (SCEs after background subtraction) (right). IR-induced SCE data shows mean ± SEM, and results from individual experiments are indicated. 120 spreads and >7,000 chromosomes per condition were analysed from three independent experiments. Knockdown of TOP3A and RAD54L was confirmed by western blotting (bottom). **(D)** HeLa cells were transfected with siCTRL, siATRX and/or siTOP3A, labelled with EdU, irradiated with 4 Gy, and then incubated with BrdU. BrdU foci were enumerated in EdU-negative G2 cells. Spontaneous foci were subtracted. All data show mean ± SEM (n = 3). Results from individual experiments, each derived from 40 cells, are indicated. **(E)** Model illustrating the regulation of HR sub-pathway usage in ATRX-proficient HeLa cells. In HeLa cells, the ATRX and RAD54L-dependent dHJ sub-pathway is predominantly used during HR. The presence of ATRX enables histone H3.3 deposition and promotes dHJ formation by counteracting TOP3A. The single loss of ATRX or H3.3 leads to failure of HR, and such a defect can be rescued upon the additional loss of TOP3A, which restores an ATRX-independent dHJ sub-pathway. *P < 0.05; **P < 0.01; ***P < 0.001; ****P < 0.0001; ns: not significant (two-tailed t test).

We next investigated if the repair pathway operating in HeLa cells in the absence of TOP3A and ATRX/H3.3 represents the dHJ sub-pathway and analyzed the formation of SCEs and BrdU repair foci. We observed that IR-induced SCE formation in ATRX-deficient HeLa cells is restored to WT levels if TOP3A is depleted (Fig. 5C). Of note, SCE formation was strictly dependent on 54L, even in the dual absence of ATRX and TOP3A (Figs. 5C). We also found that TOP3A loss in ATRX-deficient HeLa cells rescued the IR-induced BrdU repair focus formation defect (Fig. 5D). Collectively, this analysis shows that ATRX together with H3.3 antagonizes the function of TOP3A and promotes the dHJ pathway, which, we found, is strictly dependent on the function of 54L (Fig. 5E).

### HIRA promotes chromatin remodeling during SDSA

The observation that the ATRX-independent dHJ pathway, which operates in HeLa after the depletion of TOP3A, does not involve H3.3 was perhaps surprising since the chromatin remodeler HIRA can also deposit this histone variant during DSB repair^42^. Interestingly, HIRA was detected as a potential SDSA factor in our 3D screen (Fig. 1). We therefore investigated the role of HIRA during SDSA and the dHJ pathway. The depletion of HIRA did not alter γH2AX foci levels in G2-irradiated HeLa cells, consistent with the notion that it is largely dispensable for the dHJ pathway (Fig. S6A, left panel). However, loss of HIRA or H3.3 caused elevated persistent γH2AX foci in G2-irradiated U2OS cells (Fig. S6A, right panel), supporting HIRA’s potential role during SDSA. Since the formation of RAD51 foci was not affected by the depletion of HIRA (Fig. S6C), it likely functions after the strand invasion step of the SDSA pathway. We also observed that the elevated γH2AX foci level of HIRA-depleted U2OS cells is reduced to the level of WT cells after the co-depletion of TOP3A (Fig. S6A, right panel), further supporting HIRA’s role during SDSA post-strand invasion. This result was confirmed in U2OS^ATRX^ cells without Dox treatment (Fig. 6A). Upon ATRX induction by Dox treatment, we observed a reduction in γH2AX foci levels of HIRA- but not of H3.3-depleted cells, confirming that HIRA but not H3.3 is largely dispensable for the ATRX-dependent dHJ pathway (Fig. 6A). Moreover, the co-depletion of TOP3A completely rescued the elevated γH2AX foci level of Dox-treated U2OS cells devoid of H3.3 (Fig. 6A), confirming that the ATRX-independent dHJ pathway operates without H3.3. Finally, we confirmed the roles of HIRA and H3.3 during SDSA as well as the role of H3.3 during the ATRX-dependent dHJ pathway using U2OS and HeLa HR reporter cells (Fig. 6B and S6C). Collectively, our analysis shows that the histone variant H3.3 is involved in both HR sub-pathways, but the chromatin remodelers HIRA and ATRX, which are both known to deposit H3.3, exhibit specific roles during SDSA and the dHJ pathway, respectively (Fig. 6C and 5E).

**Figure 6.**
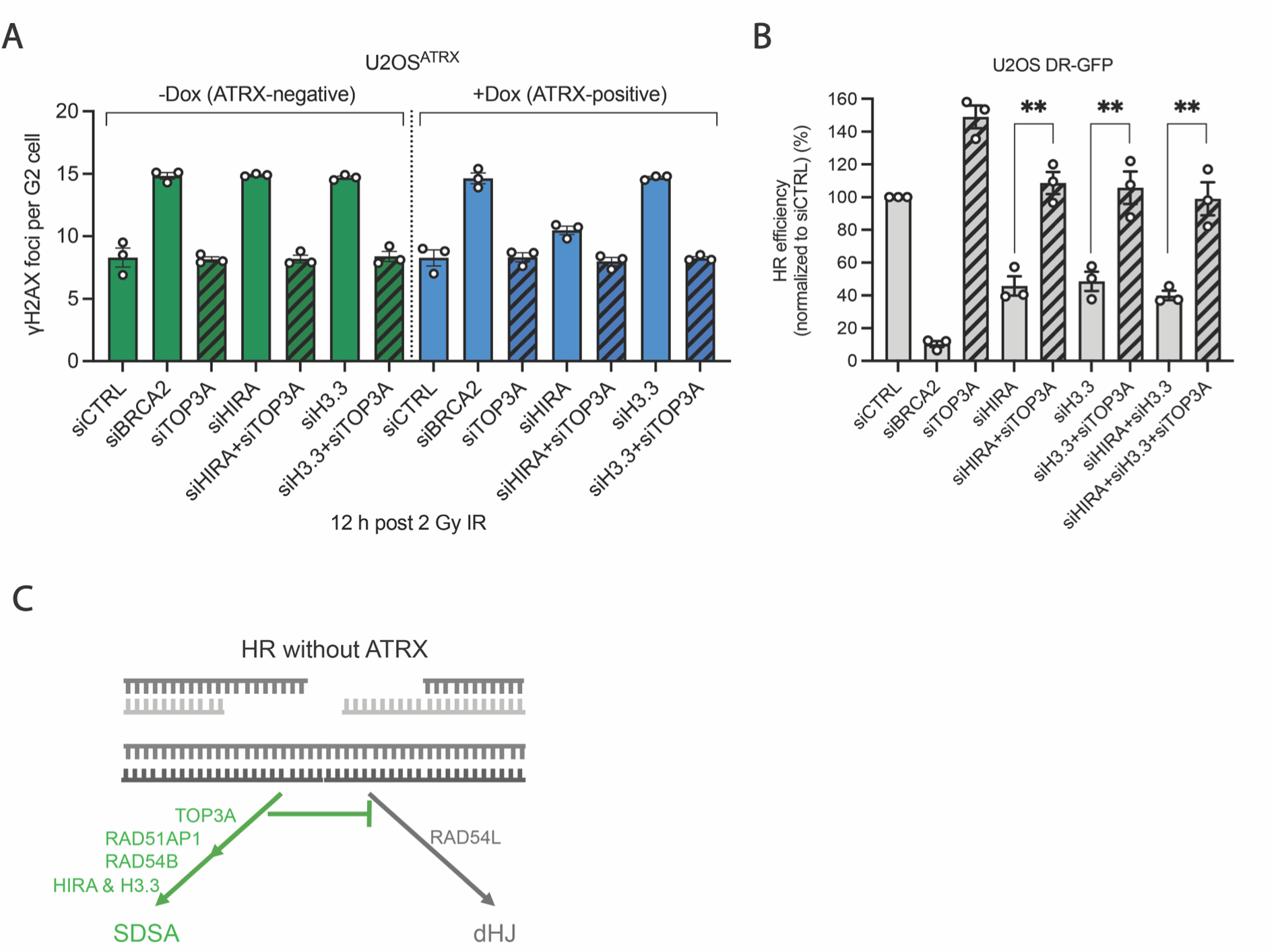
Identification of HIRA as SDSA-specific HR factor. **(A)** U2OS^ATRX^ cells, with (blue) and without (green) doxycycline-induced ATRX expression, were transfected with the indicated siRNAs, and γH2AX foci were enumerated at 12 h following 2 Gy IR in EdU-negative G2 cells. Spontaneous foci were subtracted. All data show mean ± SEM (n = 3). Results from individual experiments, each derived from 40 cells, are indicated. Knockdown was confirmed by western blotting. **(B)** U2OS DR-GFP reporter cells carry a GFP cassette that is disrupted by the I-SceI sequence and a second truncated GFP cassette. I-SceI-induced HR repair events result in the expression of functional GFP. Cells were transfected with the indicated siRNAs followed by the I-SceI plasmid transfection, and GFP-positive cells were measured by flow cytometry. All data show mean ± SEM (n = 3). Results from individual experiments are indicated. **(C)** Model illustrating the regulation of HR sub-pathway usage in intrinsically ATRX-deficient U2OS cells. In addition to RAD51AP1 and RAD54B, HIRA contributes to SDSA, but not to the dHJ sub-pathway, through its H3.3 deposition function. *P < 0.05; **P < 0.01; ***P < 0.001; ****P < 0.0001; ns: not significant (two-tailed t test).

## Discussion

Our study was designed to identify and characterize DDR pathways whose combined loss confers PARPi sensitivity. For this, we made use of the described synthetic lethality of 54L-and AP1-deficient cells^28^ and performed a PARPi 3-D CRISPR Cas9 drop-out screen using 54L KO, AP1 KO and corresponding HeLa EV cells. We utilized the Brunello library targeting 19,000 genes and confirmed the validity of the screen by identifying 54L as a sensitivity hit in AP1 KO cells and AP1 as a hit in 54L KO cells, while all three cell lines uncovered PARP1 and PARG as positive hits. Using a combinatorial evaluation of the different hits in all three cell lines, we identified factors that function specifically in the 54L pathway, specifically in the AP1 pathway or more generally confer PARPi resistance in all three cell lines. This combinatorial 3-D analysis allowed us to screen a vast number of genes, which is not easily achievable with the described dual-guide RNA screens which typically investigate combinations of factors already known to be involved in the DDR^26–27^. Moreover, dual-guide RNA screens require both gRNAs to cause gene inactivation in a single cell with multiple alleles, while our 3-D screen, utilizing a combinatorial evaluation of KO and WT cells, requires only single gene inactivation.

We then aimed to identify the pathways associated with the functions of 54L and AP1 and, based on the known roles of these factors during HR, studied in which HR sub-pathway 54L and AP1 function. For this, we applied a previously described G2-specific HR sub-pathway analysis in which we discriminate between DSB repair by SDSA and the dHJ pathway^29–30^. While SDSA is associated with short DNA repair synthesis patches and ensues without SCE formation, the dHJ pathway generates SCEs after extended DNA repair synthesis^29–30^. Importantly, this analysis revealed that 54L is involved in the dHJ pathway while AP1 operates specifically during SDSA, demonstrating that the dual loss of both HR sub-pathways is functionally equivalent to a complete loss of DSB repair by HR, similar to what is known for core HR factors. We confirmed the HR sub-pathway specificity of our identified hits by showing that 54B, which the screen identified as a factor operating in the AP1 pathway, is involved in SDSA but dispensable for the dHJ pathway. Thus, our screen identified several known as well as many hitherto undescribed HR factors, which are specifically involved in one or the other sub-pathway. The loss of an HR sub-pathway-specific factor does not significantly sensitize WT cells but confers a “BRCAness” phenotype in cells in which the other HR sub-pathway is already deficient. Since tumor cells often harbor mutations in HR sub-pathway factors^43–45^, factors of the other sub-pathway will represent targets to specifically confer an HR deficiency in the tumor while maintaining HR proficiency in the non-tumor cells. We suggest that this rationale allows the synthetic and tumor-specific generation of a “BRCAness” phenotype which holds great clinical potential.

Since 54L, 54B as well as AP1 have been described to function at the level of D-loop formation, we investigated if changes in the regulation or processing of D-loops could affect HR sub-pathway choice. Studies in yeast have suggested that two different types of D-loops can be formed during HR, one of which is preferentially processed by the BTR (yeast Sgs1-Top3-Rmi1) complex through a decatenation process^38–39^. We therefore interrogated whether TOP3A has a role in HR sub-pathway regulation and showed that loss of TOP3A leads to a shift from AP1- and 54B-dependent SDSA to the 54L-dependent dHJ pathway. This suggests that a pre-D-loop structure common to both HR sub-pathways is initially formed, which is then processed by the BTR complex into a mature D-loop compatible for SDSA. AP1 and/or 54B might exert stabilizing roles during D-loop maturation, which might be particularly important if the decatenation function of the BTR complex leads to smaller and less stable D-loops. In the absence of the BTR complex, the roles of AP1 and 54B are dispensable and the D-loop matures by the function of 54L into a structure that is compatible with the dHJ pathway. Hence, the BTR complex promotes SDSA by preventing 54L-dependent D-loop formation (see Fig. 4F).

The observation that ATRX-deficient U2OS cells switch upon depletion of TOP3A from SDSA to the dHJ pathway shows that this HR sub-pathway can operate in the absence of ATRX. We, therefore, investigated HeLa cells and show that, while the loss of ATRX confers a repair defect in this cell type, the additional loss of TOP3A allows the dHJ pathway to proceed without ATRX. We conclude from this observation that both ATRX-deficient HeLa and U2OS cells utilize the dHJ pathway in the absence of TOP3A. However, if ATRX is present, it suppresses the function of TOP3A and promotes HR repair by the dHJ sub-pathway. The observation that the role of ATRX involves the deposition of the histone variant H3.3 further suggests that this chromatin remodeling step overcomes the decatenation function of TOP3A, possibly by histone deposition within the D-loop^29^ (see Fig. 5E). This antagonizing role of ATRX and TOP3A is reminiscent of cells which employ the alternative lengthening of telomeres (ALT) pathway for telomere length maintenance, which requires either the loss of ATRX or the over-expression of TOP3A^46^.

While ATRX-dependent H3.3 deposition promotes the dHJ pathway^29^, we show in the present manuscript that SDSA involves the chromatin remodeler HIRA and also histone H3.3. While HIRA has previously been implicated in HR by depositing H3.3 at the break site^42^, our observation that it functions in concert with H3.3 specifically during SDSA suggests that distinct mechanisms of chromatin remodeling take place during SDSA and the dHJ pathway. Whereas ATRX-dependent H3.3 deposition promotes the dHJ pathway, HIRA-dependent H3.3 deposition promotes SDSA. This conclusion is supported by our CRISPR-Cas9 screen, which identified ATRX and HIRA as factors associated with 54L and AP1, respectively. One attractive model is that ATRX-dependent H3.3 deposition takes place inside the D-loop to antagonize the function of the BTR complex, whereas HIRA might deposit the histone variant H3.3 behind the D-loop to allow strand displacement once DNA repair synthesis has occurred.

In conclusion, our study reveals several novel factors involved in HR sub-pathway regulation and shows that distinct D-loop processing and chromatin remodeling processes shape the architecture of early HR intermediates, which guide HR sub-pathway choice.

## Materials and Methods

### Cell culture

U2OS (ATCC), U2OS DR-GFP, HeLa-S3 cells (ATCC), HeLa EV, HeLa RAD54L, HeLa RAD54B KO, HeLa RAD51AP1 KO, HeLa ATRX KO and HeLa pGC cells were cultured in Dulbecco’s Modified Eagle’s Medium (DMEM; Sigma), supplemented with 10% fetal bovine serum (FBS; Bio&SELL), 1% non-essential amino acid (NEAA; Sigma-Aldrich) and 1% penicillin/streptomycin (Sigma-Aldrich). Genetically modified U2OS cells with doxycycline-inducible ATRX expression (a kind gift from David Clynes) were maintained in DMEM (Sigma-Aldrich), supplemented with 10% tetracycline-free FBS (Takara), 1.2% L-Glutamine (Sigma-Aldrich) and 1% penicillin/streptomycin (Sigma-Aldrich). All cell lines were grown at 37°C in humidified incubators containing 5% CO_2_.

### Generation of CRISPR KO cell lines

HeLa ATRX KO, RAD54L KO and RAD51AP1 KO cells were generated by transfecting HeLa cells with pX459v2 vectors (Addgene #62988) carrying the sgRNA targeting sequences using FUGENE HD (Promega) according to manufacturer’s instructions. 48 h after transfection, cells were selected with puromycin (1 μg/ml) or blasticidin (7.5 μg/ml) for 10 days, followed by reseeding for clonal expansion. The selected clones were verified by western blotting and genomic DNA sequencing. The sgRNA oligonucleotides used for generating the knockouts can be found in Table 1.

**Table 1:**
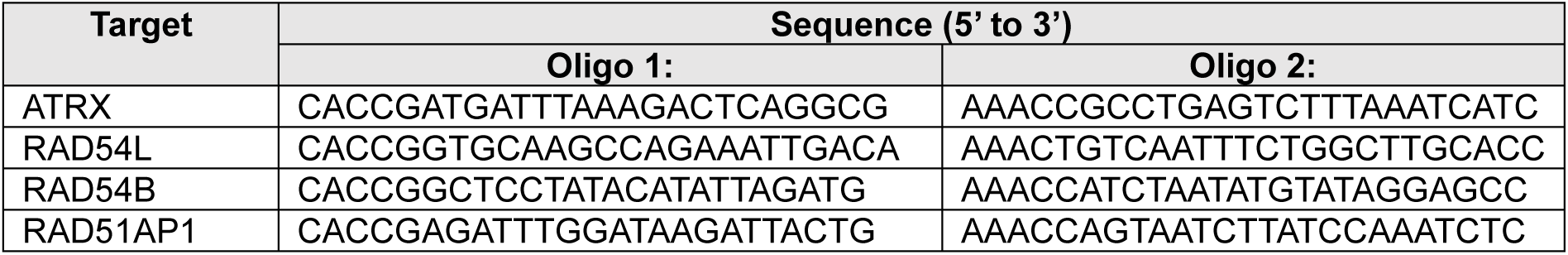
sgRNAs used in this study.

### S2 CRISPR-Cas9-knockout Screen

For each cell line, 3.64 × 10^6^ cells were transduced with the Brunello lentiviral library at an MOI of ∼0.3 in the presence of 8 µg/ml polybrene to obtain a 500x coverage, and cells were selected for 4 days with 1.5 μM of Puromycin. Post selection, 60 × 10^6^ cells were treated with 5 µM Olaparib, and 60 × 10^6^ cells were left untreated. Cells were collected for DNA extraction at day 0 and day 14.

### Library preparation

Genomic DNA (gDNA) was extracted from the frozen cell pellets using the QIAamp^®^ DNA Blood Maxi Kit (QIAGEN) according to the manufacturer’s instructions. Polymerase chain reaction (PCR) was performed as described^47^.PCR products were cleaned using a two-step AMPure XP bead purification (Beckman Coulter) protocol, and quality control (QC) was performed using the TapeStation D1000 Kit (Agilent), both according to the manufacturer’s instructions.

### Illumina sequencing

Processed PCR products were pooled and sent frozen to the Genomics Core Facility of the Institute of Molecular Biology (IMB) in Mainz. There, the pool of libraries was sequenced as 126 nt single-end reads on an Illumina NextSeq 2000 platform using two P3 (100-cycles) flow cells. Sequencing yielded in 56 to 517 million reads per sample.

### Bioinformatic processing and analysis of CRISPR-Cas9 screen sequencing data

#### Read QC

Raw sequencing reads (FASTQ files) were assessed using FastQC (https://www.bioinformatics.babraham.ac.uk/projects/fastqc/) to evaluate per-base quality profiles, adapter content, and sequence duplication levels. FastQC reports were reviewed to identify potential sequencing artifacts (e.g., low quality reads or base composition bias).

#### sgRNA quantification and screen-level quality metrics

sgRNA read counting was performed using the MAGeCK count module, using the reference sgRNA library annotation as input. Reads were assigned to sgRNAs by matching the guide sequence within each read to the expected protospacer sequence. Standard screen-level QC metrics were computed from the MAGeCK count output, including total reads, fraction of reads mapped to the sgRNA library, distribution of sgRNA counts, sgRNAs with zero reads, Gini-index and negative selection quality control.

#### Exploratory sample relationships

To visualize overall sample similarity and replicate concordance, normalized sgRNA count matrices were transformed (log2) and used for principal component analysis (PCA). PCA was performed at the sgRNA level after normalization by the MAGeCK count module to reduce differences in sequencing depth across samples.

#### Gene-level effect estimation using MAGeCK-MLE and MAGeCKFlute

Gene-level selection effects were estimated using MAGeCK-MLE (Maximum Likelihood Estimation using a generalized linear model), a part of MAGeCK-VISPR^48^, with a design matrix encoding baseline (T0) and condition-specific coefficients for each treatment and cell line. In brief, the model estimates per-gene effect sizes (“beta scores”) reflecting the direction and magnitude of enrichment/depletion associated with each condition relative to baseline.

Normalization was performed prior to model fitting using median normalization anchored to a defined set of non-essential control genes, as implemented in MAGeCK-MLE. To enable comparisons across conditions with potentially different growth dynamics, beta scores were additionally adjusted using the MAGeCKFlute “cell_cycle” normalization step, which uses the behavior of essential genes to account for proliferation-related shifts across samples. Effect sizes (beta-derived metrics) were used for prioritization and comparisons across conditions.

#### Derivation of treatment-specific effects and between-cell line comparisons

Within each cell line, treatment-specific effects were computed by contrasting the treatment coefficient with the matched non-treated control coefficient, both referenced to the same baseline. Afterwards, treatment-specific delta-betas were calculated as:

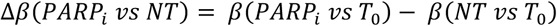

Between-cell line comparisons were performed using these delta-beta values, following normalization as described above.

#### Visualization and intersection analysis of candidate dependencies (revised)

Downstream analyses were performed in Python using pandas, NumPy, matplotlib, and adjustText. Pairwise scatter plots were generated by plotting delta-beta values of one cell line against another (x- and y-axes, respectively). To delineate effect regions in an unbiased manner, cutoff lines were defined empirically from the distribution of values in each comparison. The cutoff lines were centered around 0 as done by the MAGeCKFlute R package, but defined over two standard deviations as follows:

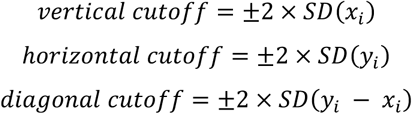

To compare gene sets across multiple pairwise cell line comparisons, genes were first partitioned into region-specific lists based on their position relative to the vertical, horizontal, and diagonal cutoff boundaries described above. For each pairwise scatter plot, region membership was used to generate discrete gene sets corresponding to specific effect patterns. These region-derived gene sets from the different cell line comparisons were then compared using an UpSet plot (Python upsetplot; visualization with matplotlib and matplotlib-venn) to summarize intersections. Intersections were subsequently used to group genes into analysis categories (SDSA-associated, dHJ-associated, HR-core factors, common regulators of SDSA and dHJ) for downstream reporting and visualization.

#### Annotated scatter plots of intersection-defined gene groups (updated)

To visualize the intersection-defined gene groups, we regenerated the three pairwise delta-beta scatter plots and overlaid group assignments derived from the UpSet intersections. Genes belonging to a given intersection group were colored using the same palette as in the UpSet plot to maintain consistent group identity across visualizations, while all other genes were shown in a neutral background color.

For each pairwise comparison, we additionally labeled a subset of “top” genes per intersection group based on axis-combination metrics computed from the plotted coordinates. Specifically, for each gene with coordinates (x, y) (delta-beta in the two compared cell lines), we calculated (i) the difference between axes, d = y − x, and (ii) the sum of axes, s = x + y. Genes with the most negative or positive difference d were labelled in the group of dHJ-associated and SDSA-associated genes, respectively. In the groups of common regulators and HR-core factors, genes with the most negative sum s were selected for labelling. Gene labels were placed using adjustText to reduce overlap.

### RNA interference

Cells were transfected with 30 or 50 nM siRNA using Lipofectamine RNAiMAX (Invitrogen) according to manufacturer’s instructions. The siRNA sequences used can be found in Table 2. Cells were typically collected for immunofluorescence and western blotting 48 h post siRNA transfection.

**Table 2:**
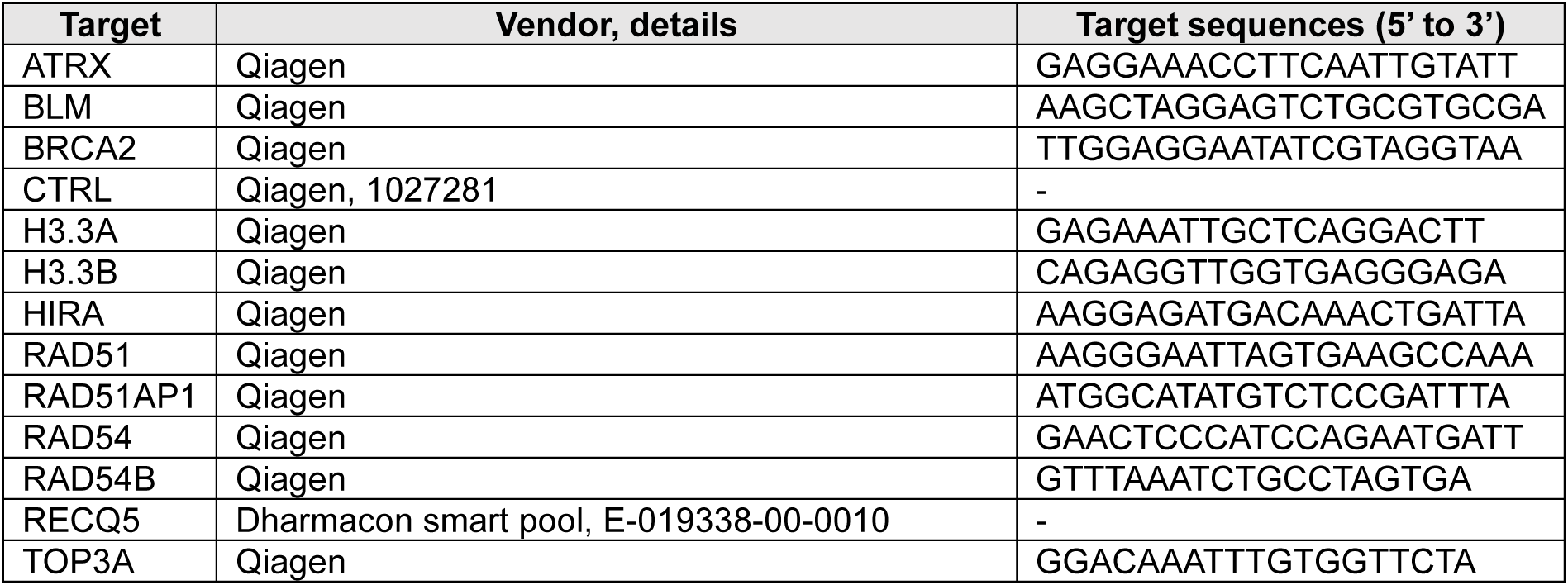
siRNA used in this study.

### Western blotting

Knockdown and knockout efficiencies were confirmed by western blotting. Cells were collected by trypsinization and lysed with Cell Signaling lysis buffer (Cell Signaling, #9803) supplemented with 1x PhosSTOP (Roche) and 1x cOmplete EDTA-free protease inhibitor cocktail (Roche), followed by sonication. Protein extracts were boiled with Laemmli buffer (10% sodium dodecyl sulfate, 300 mM Tris-HCl, 10 mM β-mercaptoethanol, 50% glycine, 0.02% bromophenol blue) at 95°C for 5 min. Proteins were separated by SDS-PAGE and transferred to 0.2 µm nitrocellulose membranes (Amersham) using wet transfer. After the transfer, membranes were blocked with 5% skimmed milk in TBS/0-1% Tween-20 (TBS-T) at room temperature and incubated with the relevant primary antibodies overnight at 4°C. Afterwards, the membranes were washed with TBS-T and incubated with horseradish peroxidase (HRP)-conjugated secondary antibodies for 1 h at room temperature. After the membranes were washed with TBS-T, the immunoblots were developed using Western Blot Quantum or Sirius kits (Advansta) and the signals were detected using the Fusion FX system (Vilber Lourmat). Primary and secondary antibodies used for western blotting can be found in Table 3.

**Table 3:**
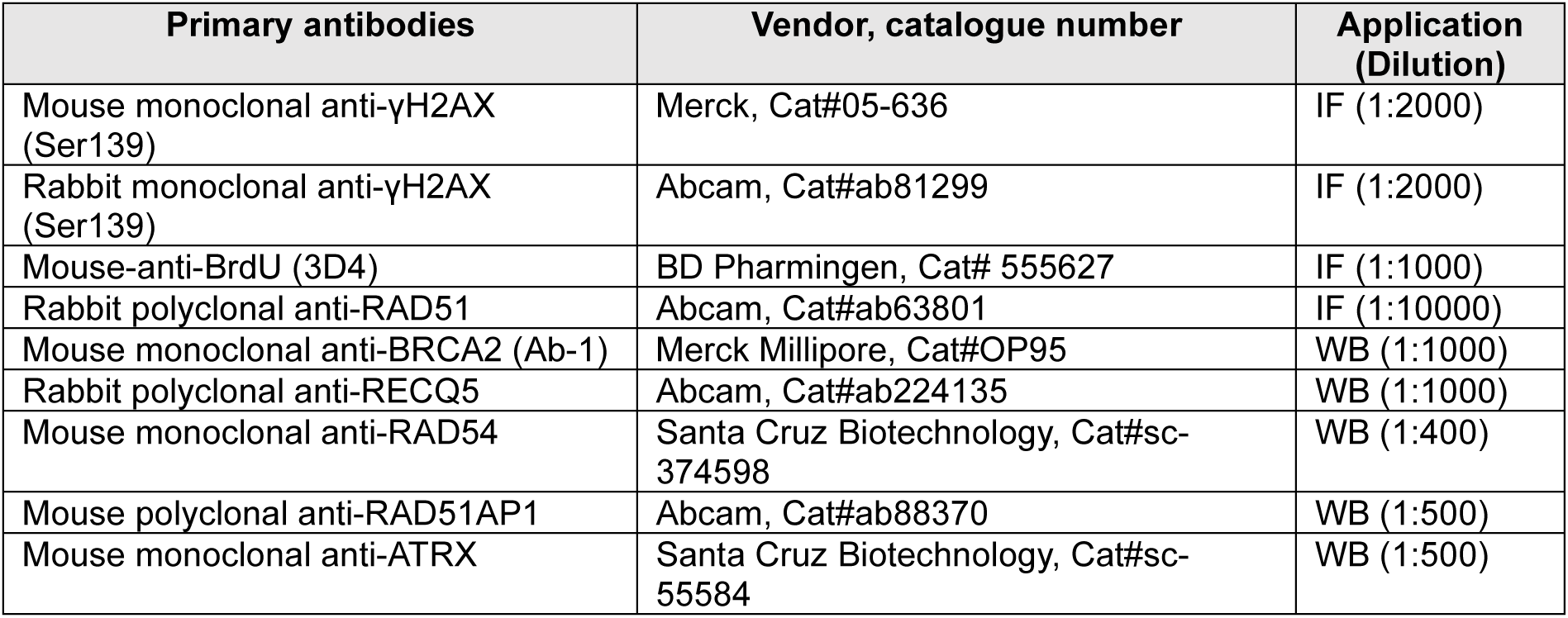
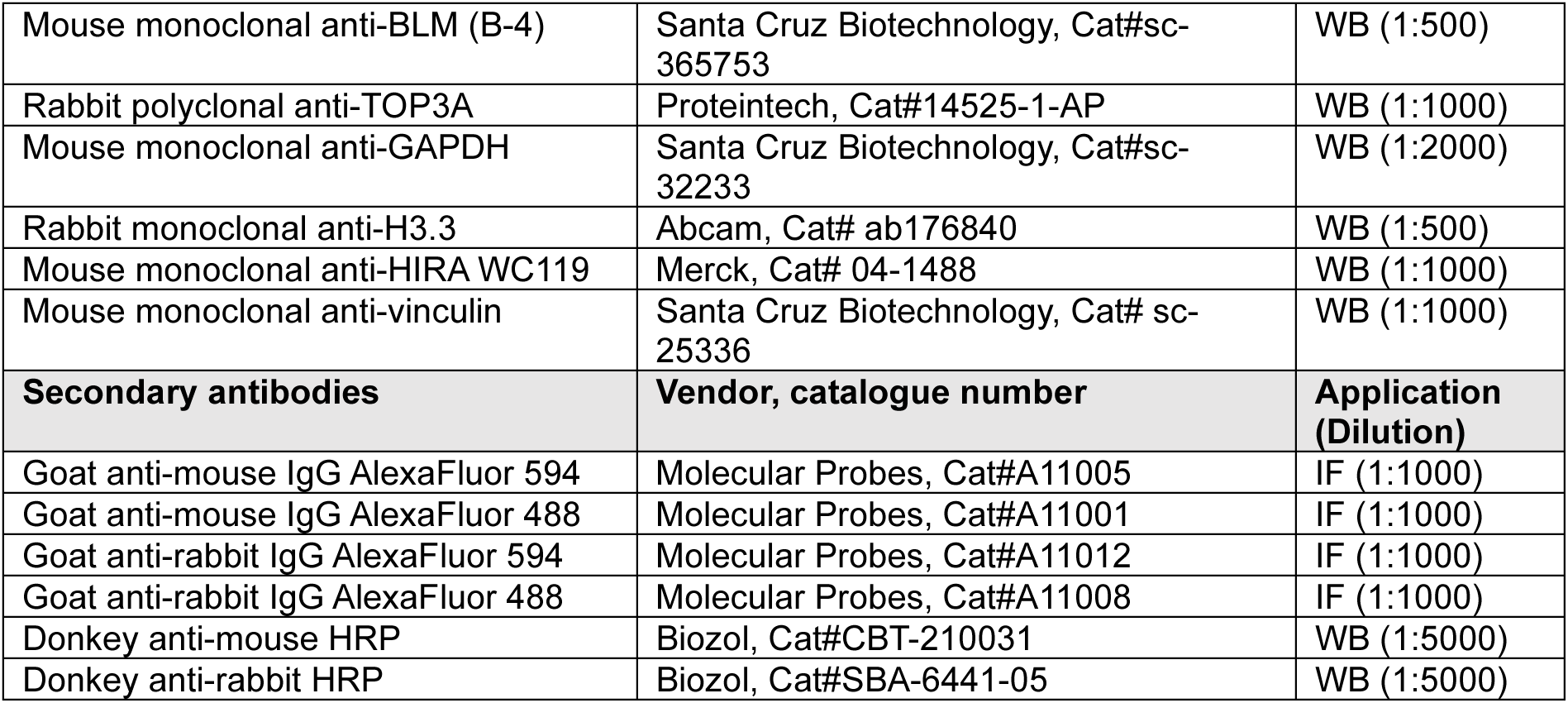
Antibodies used in this study.

### Irradiation and drug treatment

For X-ray irradiation treatment, cells were irradiated at 90 kV and 19 mA or 33.7 mA for 2 or 4 Gy with an aluminum filter using an X-ray generator (GE Inspection Technologies). The doses and timing of olaparib (Selleck) used are indicated in the respective figures.

### Cell cycle-specific DSB repair analyses

The cell cycle-specific DSB repair analyses were carried out as described previously^29^. For analyses in response to IR treatment, the thymidine analog EdU (10 μM; Baseclick, BCN-001) was added to the cells 30 min before irradiation and maintained throughout the experiment. For analyses in response to olaparib treatment, 10 μM EdU was added 1 h before harvesting. After cell fixation and immunostaining, DAPI and EdU intensities were measured and plotted in a diagram (EdU signal against DAPI signal) using a Zeiss microscope and MetaCyte software (Metasystems). EdU-positive cells were categorized as S-phase cells, whereas the EdU-negative cells were identified as G1 or G2 cells based on their DAPI intensity (corresponding to their DNA content). To analyze DSB repair in G2 phase cells, all EdU-positive cells were excluded in this study, and only EdU-negative cells with high DAPI intensity were selected for evaluation.

### Immunofluorescence

Cells cultured on coverslips were washed with PBS, fixed with 2% paraformaldehyde (PFA), washed again with PBS and permeabilized with 0.2% Triton X-100 in PBS for 10 min, followed by washing with 0.1% Triton X-100 in PBS. Afterwards, the cells were blocked with 1x Roti Immunoblock (Carl Roth) for 1 h at 4°C and incubated with primary antibodies diluted in 1x Roti Immunoblock overnight at 4°C. Cells were washed with 0.1% Triton X-100 in PBS, incubated with secondary antibodies diluted in 1x Roti Immunoblock for 1 h and washed with 0.1% Triton X-100 in PBS. To stain the EdU for cell-cycle specific analyses, cells were incubated with EdU Click-iT Kit (Baseclick) for 30 min according to the manufacturer’s instructions. Followed by washing with 0.1% Triton X-100 in PBS, cell nuclei were stained with 4’,6-diamidino-2-phenylindole (DAPI) for 10 min at room temperature. After the DAPI staining, cells were mounted with Vectashield antifade mounting media (Vector Laboratories). All immunofluorescence analyses were performed using a Zeiss microscope with Metafer4 software (MetaSystems). Primary and secondary antibodies used for immunofluorescence can be found in Table 3.

Representative images were taken using a Zeiss microscope with ApoTome.2 (Zeiss) and the Metafer4 (MetaSystems) and the Zen 3.0 (Zeiss) software. They were then processed using the ImageJ software, where brightness and contrast were adjusted for every image if necessary.

### BrdU incorporation assay

The assay was performed similarly as previously described^29^. In brief, cells were grown on coverslips and transfected with siRNA. Following 48 h siRNA treatment, cells were labelled with 10 μM EdU for 1 h and irradiated with 4 Gy. Cells were then immediately washed with PBS, incubated with 50 μM BrdU for additional 8 or 10 h and fixed at the indicated time points with 3% PFA. DNA was then denatured using 2.5 M HCl, followed by extensive washing with PBS. Afterwards, the cells were permeabilized with 0.5% Triton X-100 in PBS and washed with PBS-0.1% Triton X-100. The cells were then blocked with 5% FCS in PBS-0.1% Triton X-100. Staining was performed as described in immunofluorescence (except the antibodies were diluted in 5% FCS in PBS-0.1% Triton X-100). BrdU foci were enumerated manually in EdU-negative G2 phase cells.

### Sister chromatid exchanges

HeLa cells were transfected with siRNA and incubated in media containing 10 μM BrdU for 48 h. HeLa cells were then irradiated with 2 Gy and incubated for another 8 h before harvesting. To enrich mitotic cells and override the DNA damage checkpoint induced by IR, irradiated HeLa cells were treated with 1 µg/ml colcemid and 5 nM caffeine 3 h before harvesting. Undamaged HeLa cells were treated with 1 µg/ml colcemid for 3 h after 48 h BrdU incubation. For U2OS^ATRX^ cells, cells were incubated with 10 μM BrdU for 48 h. Irradiated U2OS^ATRX^ cells were incubated for further 16 h and treated with 50 nM calyculin A (LC Laboratories) for 1 h before harvesting. Cells were harvested, swollen in 75 mM KCl for 37°C and fixed with ice-cold methanol:acetic acid (3:1). The fixed cells containing the chromosomes were spread on glass slides and air-dried overnight. The dried glass slides were then incubated with 5 µg/mL Hoechst 33258, washed with water, and covered with an irradiation buffer (200 mM Na2HPO4 + 4 mM citric acid, pH 7.1-7.2) for exposure to UV light at 9 joules/cm² for 45 min. After the UV treatment, slides were incubated in 2x SSC buffer at 55°C and stained with 6% Giemsa. Images of chromosome spreads were captured using a Zeiss microscope with Metafer4 software (MetaSystems).

### HR reporter assays

The efficiency of HR repair was measured in U2OS DR-GFP cells (a gift from Jeremy Stark) or HeLa pGC cells (a gift from Jochen Dahm-Daphi). Cells were first seeded in 24-well plates and transfected with siRNA using Lipofectamine RNAiMAX (Invitrogen). One day after siRNA transfection, the cells were transfected with either 0.75 μg empty control plasmid (pUC19) or I-SceI endonuclease expression plasmid (pBL464-pCBASce) using Lipofectamine LTX (Invitrogen) according to the manufacturer’s instructions, followed by an incubation for another 24-72 h. The cells were then collected, and the percentage of GFP-positive cells was measured using an S3 cell sorter (Biorad).

### Survival assays

Cell survival in response to different doses of olaparib was tested by performing a CellTiterGlo® 2.0 Viability assay or a growth curve analysis.

For the CellTiter-Glo® 2.0 Viability assay, cells were seeded in 35 mm dishes and transfected with siRNA the next day. One day after transfection, 100 or 250 cells per well were reseeded in triplicate in white 96-well plates. Treatment with DMSO, 5 µM or 10 µM olaparib started 24 h after reseeding, and cell survival was measured five days later using the CellTiter-Glo® 2.0 assay (Promega) according to the manufacturer’s instructions. Luminescence signals were read using a Spark multimode microplate reader (Tecan). The signals obtained from cells treated with olaparib were normalized to those of untreated cells.

For growth curve analyses, cells were seeded in 60 mm dishes and transfected with siRNAs. 24 h later, the cells were reseeded in duplicate in 6-well plates, and treatment with DMSO or olaparib started. The cell number was determined using Neubauer counting chambers on days four and seven. On day four, olaparib-treatment was renewed. The absolute cell numbers were normalized to DMSO-treated samples.

### Quantitative RT-PCR

Analysis of RAD54B expression after siRNA depletion was performed using quantitative PCR and reverse transcription. 48 h after siRNA transfection, cells were harvested and RNA extraction was performed using MasterPure Complete DNA and RNA Purification Kit (Lucigen) according to the manufacturer’s instructions. 1 µg RNA was transcribed into DNA using RevertAid FirstStrand cDNA reverse transcription kit (Thermo Fisher) according to the manufacturer’s instructions. The generated cDNA was analyzed by quantitative PCR using FastStart Universal SYBR Green Master mix (Roche) according to the manufacturer’s instructions, and primers targeting RAD54B and GAPDH as a housekeeping gene (RAD54B: fwd 5’ gcttgactgtgagtgtacagg 3’, rev: 5’ ctggtgatgtggaccaag 3’; GAPDH: fwd: 5’ ggtgaaggtcggagtcaacg 3’, rev: 5’ gtagttgaggtcaatgaaggggtc 3’). Amplification was performed using the Comparative CT settings of a StepOne Real-Time PCR System (Applied Biosystems) with the following programme: 95°C for 10 min, 40 cycles of 95°C for 10 s and 60°C for 30 s, followed by a melting curve analysis to confirm primer specificity. All samples were run in triplicate, and the Ct value was normalized to GAPDH and compared to siCTRL samples.

#### Quantification and statistical analysis

Unless otherwise specified in the figure legends, all data were derived from at least n = 3 replicates and for each experiment at least 40 nuclei or 40 chromosome spreads were analyzed. Background foci were subtracted from the mean values. Column plots show the mean value and the error bars show the SEM between the experiments. P values were obtained by two-tailed t-test and compare the mean values of independent experiments (*P < 0.05; **P < 0.01; ***P < 0.001; ****P < 0.0001). All statistical analyses were performed using Prism 11 software (GraphPad).

## Declarations

### Data Availability

Supplemental Information accompanies this paper. The datasets of this article are either included within the article and its additional files, or are available upon request.

## Acknowledgments

We thank David Clynes, Jeremy Stark and Jochen Dahm-Daphi for providing cell lines, IMB Core Facilities, Amelie Fischer and Andrés Cruz-García for experimental help and pilot studies, Cornelia Schmitt and Bettina Basso for excellent technical support and Wolf D. Heyer for the helpful discussions. This work was funded by the Federal Ministry of Research, Technology and Space (02NUK034A and 02NUK054C), the Deutsche Forschungsgemeinschaft (DFG, German Research Foundation) – Project-ID 393547839 – SFB 1361.

## Author contributions

K.C.C., A.K., J.E. and B.S. performed experiments. K.C.C., A.K., J.E., B.S. and A.B. interpreted data. R.N.C., B.S. and J.E. established knockout cell lines. M.L. conceived experiments and wrote the paper, aided by K.C.C., A.K, J.E, B.S. and A.B.

---

**Figure S1.**
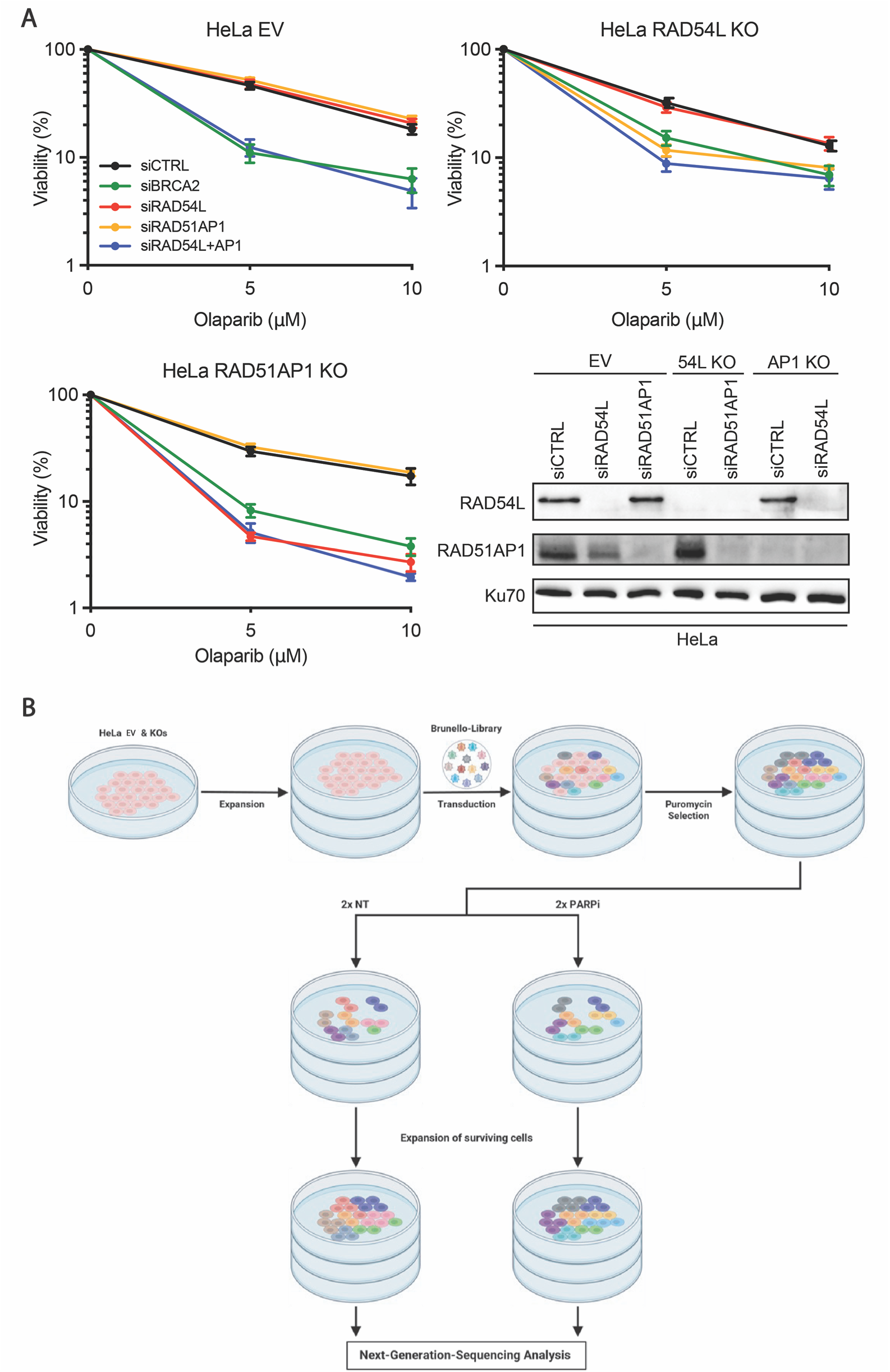
(Related to Figure 1: CRISPR screens identify synthetic lethal targets in HeLa EV, RAD54L KO and RAD51AP1 KO cells). **(A)** Survival of HeLa EV, RAD54L and RAD51AP1 KO cells transfected with the indicated siRNAs upon treatment with 5 and 10 µM of olaparib or untreated. Cell viability was measured using CellTiter-Glo assay. All data show mean ± SEM (n=3). **(B)** Schematic of genome-wide CRISPR/Cas9 screening workflow.

**Figure S2.**
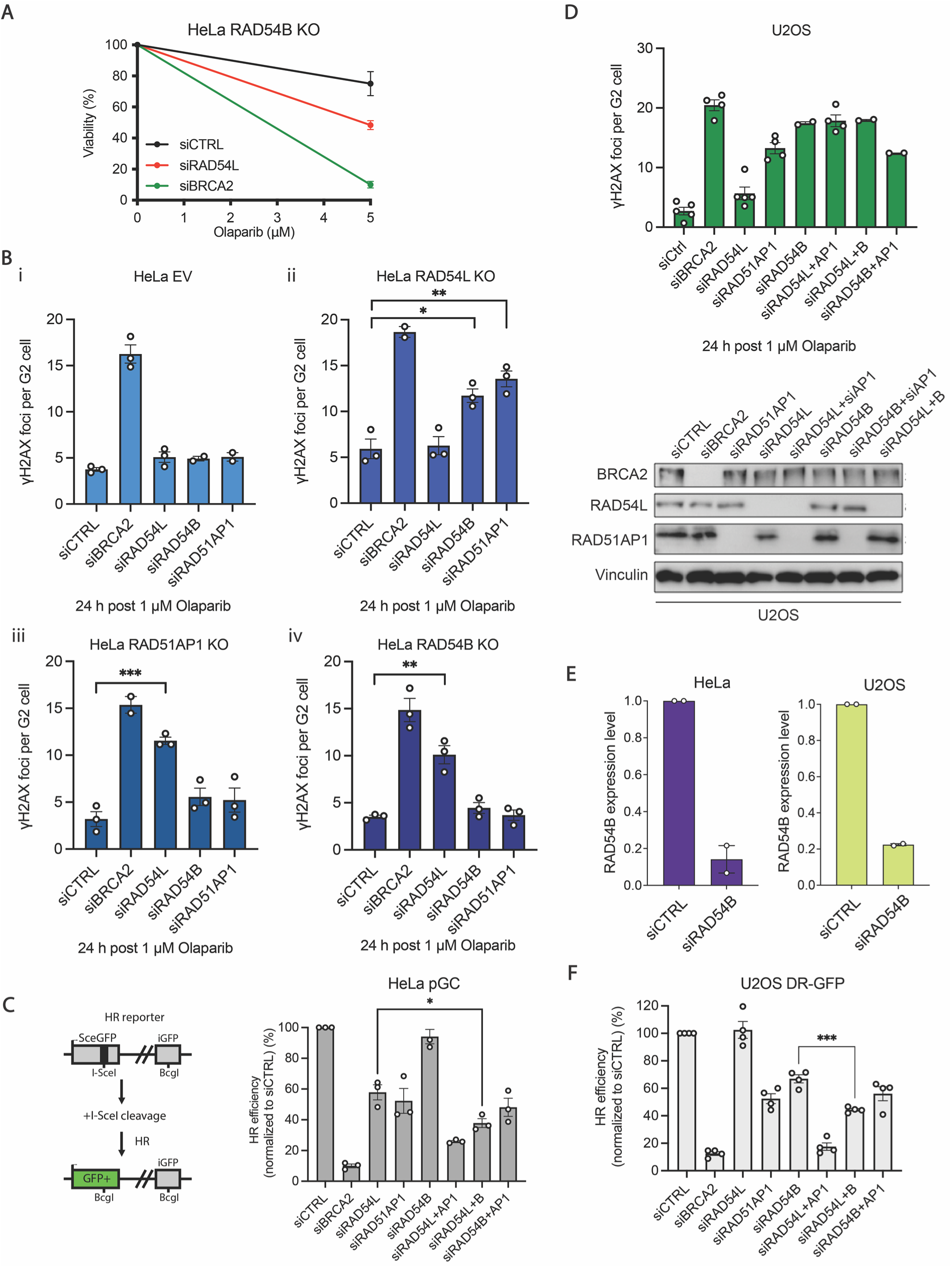
(Related to Figure 2: RAD51AP1/RAD54B and RAD54L define distinct sub-pathways of HR). **(A)** Survival of HeLa RAD54B KO cells transfected with the siCTRL, siRAD54L or siBRCA2 upon treatment with 5 µM of olaparib or untreated. Cell viability was measured using growth curve analyses. All data show mean ± SEM (n=3). **(B)** HeLa (i) EV, (ii) RAD54L KO, (iii) RAD51AP1 KO and (iv) RAD54B KO cells were transfected with the indicated siRNAs, and γH2AX foci were enumerated after treatment with 1 µM olaparib for 24 h in EdU-negative G2 cells. Spontaneous foci were subtracted. All data show mean ± SEM (n = 2-3). Results from individual experiments, each derived from 40 cells, are indicated. **(C)** HeLa pGC reporter cells carry a GFP cassette that is disrupted by the I-SceI sequence and a second truncated GFP cassette (left). I-SceI-induced HR repair events result in the expression of functional GFP. HeLa pGC cells were transfected with the indicated siRNAs, followed by transfection with the I-SceI expression plasmid to induce site-specific DSBs. The percentage of GFP-positive cells was measured by flow cytometry. All data show mean ± SEM (n=3). All data show mean ± SEM (n = 3). Results from individual experiments are indicated. **(D)** (Top) U2OS cells were transfected with the indicated siRNAs, and γH2AX foci were enumerated after treatment with 1 µM olaparib for 24 h in EdU-negative G2 cells. Spontaneous foci were subtracted. All data show mean ± SEM (n = 2-5). Results from individual experiments, each derived from 40 cells, are indicated. (Bottom) Knockdown of BRCA2, RAD51AP1 and RAD54L was confirmed by western blotting. **(E)** Quantitative RT–PCR analysis of RAD54B mRNA expression levels in HeLa and U2OS cells after transfection with siCTRL or siRAD54B. Expression levels were normalized to GAPDH, with siCTRL defined as 100%. All data show mean ± SEM (n = 2). **(F)** U2OS DR-GFP cells were transfected with the indicated siRNAs, followed by transfection with the I-SceI expression plasmid to induce site-specific DSBs. The percentage of GFP-positive cells, representing HR repair events, was measured by flow cytometry. All data show mean ± SEM (n = 4). Results from individual experiments are indicated. *P < 0.05; **P < 0.01; ***P < 0.001; ****P < 0.0001; ns: not significant (two-tailed t test).

**Figure S3.**
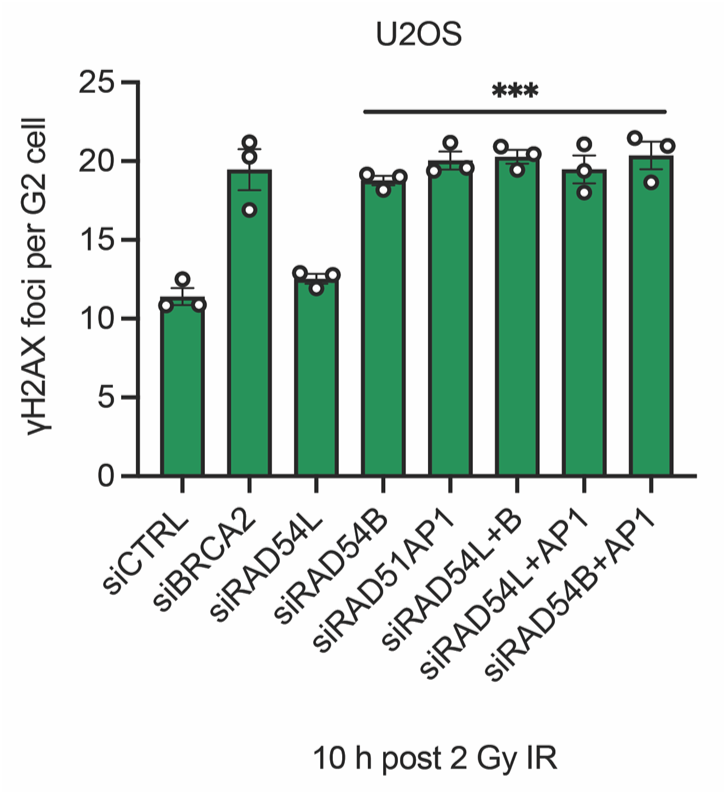
(Related to Figure 3: ATRX induction triggers a switch from RAD51AP1/RAD54B-dependent SDSA to a RAD54L-dependent dHJ sub-pathway). U2OS cells were transfected with the indicated siRNAs, and γH2AX foci were enumerated 10 h following 2 Gy IR in EdU-negative G2 cells. Spontaneous foci were subtracted. All data show mean ± SEM (n = 3). Results from individual experiments, each derived from 40 cells, are indicated. *P < 0.05; **P < 0.01; ***P < 0.001; ****P < 0.0001; ns: not significant (two-tailed t test).

**Figure S4.**
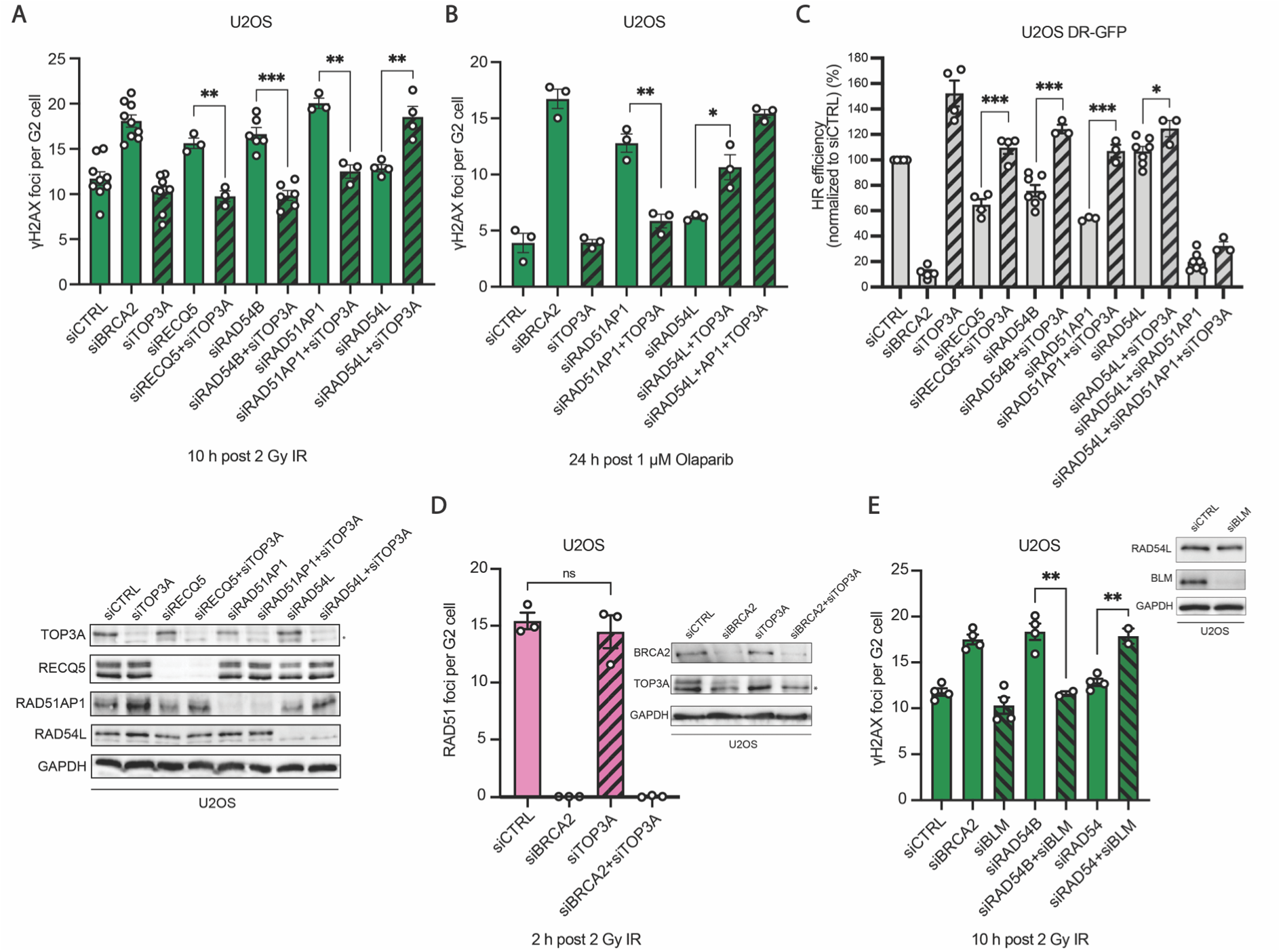
(Related to Figure 4: Loss of TOP3A function in ATRX-deficient U2OS cells causes a shift in HR sub-pathway usage from SDSA to the RAD54L-dependent dHJ sub-pathway). **(A)** (Top) U2OS cells were transfected with the indicated siRNAs, and γH2AX foci were enumerated 10 h following 2 Gy IR in EdU-negative G2 cells. Spontaneous foci were subtracted. All data show mean ± SEM (n = 3-9). Results from individual experiments, each derived from 40 cells, are indicated. (Bottom) Knockdown of TOP3A, RECQ5, RAD51AP1 and RAD54L was confirmed by western blotting. **(B)** U2OS cells were transfected with the indicated siRNAs, and γH2AX foci were enumerated after treatment with 1 µM olaparib for 24 h in EdU-negative G2 cells. Spontaneous foci were subtracted. All data show mean ± SEM (n = 3). Results from individual experiments, each derived from 40 cells, are indicated. **(C)** U2OS DR-GFP cells were transfected with the indicated siRNAs, followed by transfection with the I-SceI expression plasmid to induce site-specific DSBs. The percentage of GFP-positive cells, representing HR repair events, was measured by flow cytometry. All data show mean ± SEM (n = 3-7). Results from individual experiments are indicated. **(D)** U2OS cells were transfected with siCTRL, siBRCA2 and siTOP3A, irradiated, and RAD51 foci were enumerated 2 h following 2 Gy IR in EdU-negative G2 cells (left). Spontaneous foci were subtracted. All data show mean ± SEM (n = 3). Results from individual experiments, each derived from 40 cells, are indicated. Knockdown of BRCA2 and TOP3A was confirmed by western blotting (right). **(E)** U2OS cells were transfected with the indicated siRNAs, and γH2AX foci were enumerated 10 h following 2 Gy IR in EdU-negative G2 cells (left). Spontaneous foci were subtracted. All data show mean ± SEM (n = 2-4). Results from individual experiments, each derived from 40 cells, are indicated. Knockdown of BLM was confirmed by western blotting (right). *P < 0.05; **P < 0.01; ***P < 0.001; ns: not significant (two-tailed t test).

**Figure S5.**
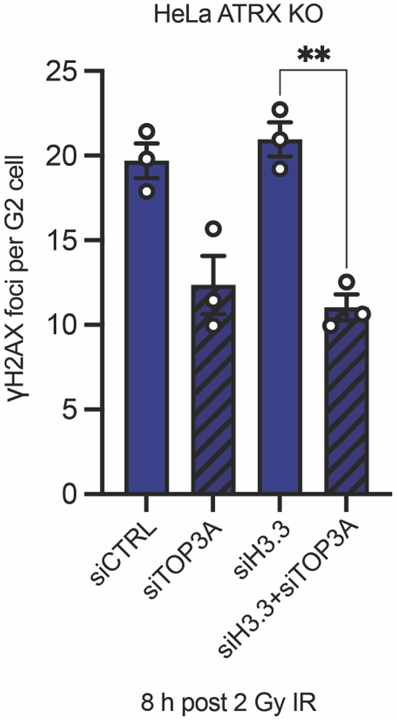
(Related to Figure 5: ATRX, together with H3.3, antagonizes TOP3A function and promotes the dHJ pathway in HeLa cells). HeLa ATRX KO cells were transfected with the indicated siRNAs, and γH2AX foci were enumerated 8 h following 2 Gy IR in EdU-negative G2 cells. Spontaneous foci were subtracted. All data show mean ± SEM (n = 3). Results from individual experiments, each derived from 40 cells, are indicated. The CRISPR-based ATRX KO cell line was tested by sequencing and western blotting. Knockdown of TOP3A and H3.3 was confirmed by western blotting. *P < 0.05; **P < 0.01; ***P < 0.001; ns: not significant (two-tailed t test).

**Figure S6.**
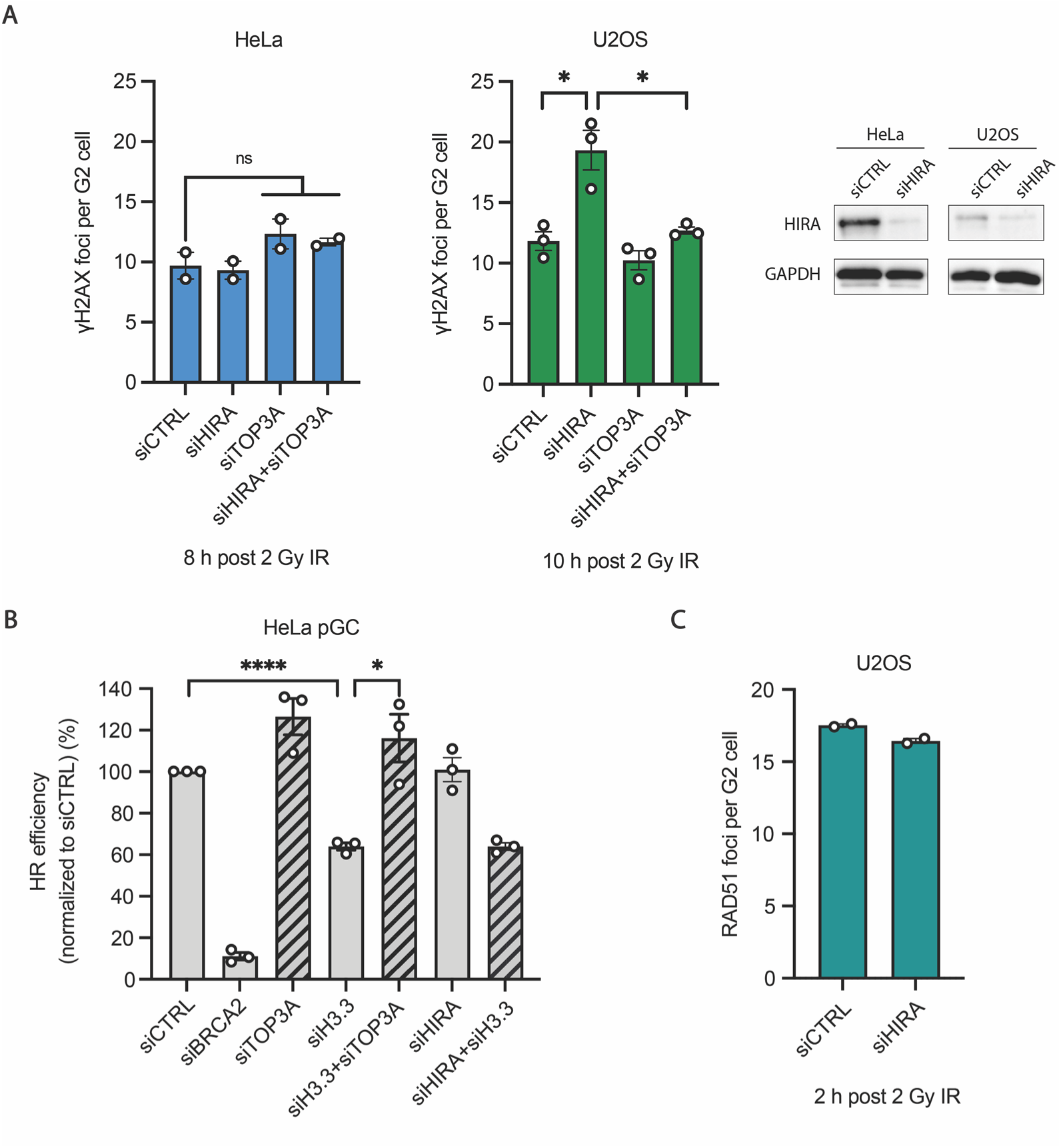
(Related to Figure 6: Identification of HIRA as SDSA-specific HR factor). **(A)** HeLa and U2OS cells were transfected with the indicated siRNAs, irradiated with 2 Gy, and γH2AX foci were enumerated at 8 h (HeLa) or 10 h (U2OS) post-irradiation in EdU-negative G2 cells. Spontaneous foci were subtracted. All data show mean ± SEM (n = 2-3). Results from individual experiments, each derived from 40 cells, are indicated. Knockdown of HIRA was confirmed by western blotting (right). **(B)** HeLa pGC cells were transfected with the indicated siRNAs, followed by transfection with the I-SceI expression plasmid to induce site-specific DSBs. The percentage of GFP-positive cells, representing HR repair events, was measured by flow cytometry. All data show mean ± SEM (n = 3). Results from individual experiments are indicated. **(C)** U2OS cells were transfected with the siHIRA and RAD51 foci were enumerated 2 h following 2 Gy IR in EdU-negative G2 cells. Spontaneous foci were subtracted. All data show mean ± SEM (n = 2). Results from individual experiments, each derived from 40 cells, are indicated. *P < 0.05; **P < 0.01; ***P < 0.001; ns: not significant (two-tailed t test).

